# Oncogenic Ras activation in permissive somatic cells triggers rapid onset phenotypic plasticity and elicits a tumour-promoting neutrophil response

**DOI:** 10.1101/2023.11.10.566547

**Authors:** Abigail M. Elliot, Isabel Ribeiro Bravo, Jeanette Astorga Johansson, Esme Hutton, Richard Cunningham, Henna Myllymäki, Kai Yee Chang, Justyna Cholewa-Waclaw, Yiyi Zhao, Mariana Beltran, Ross Dobie, Amy Lewis, Philip M. Elks, Carsten Gram Hansen, Neil Henderson, Yi Feng

## Abstract

Oncogenic driver mutations are frequently found in normal tissues, suggesting that additional non-genetic factors are required for tumourigenesis. Phenotypic plasticity is an important gateway to malignancy and inflammation can fuel tumourigenesis, however, little is known about when and how these hallmarks first arise. Using single-cell transcriptomics and *in vivo* live imaging we have characterised the immediate cell intrinsic and innate immune responses during the first 24 hours following oncogenic Ras activation, in an inducible zebrafish model of HRAS^G12V^-mediated skin tumour initiation. We found that only a subset of basal keratinocytes, but not superficial keratinocytes, are susceptible to RAS-driven phenotypic plasticity. These preneoplastic cells undergo dedifferentiation and partial EMT, resembling malignant cells observed in human squamous cell carcinoma (SCC). Strikingly, the same subset instigates the development of tumour-promoting neutrophils, which in turn enhance preneoplastic cell proliferation. Our findings demonstrate that the effects of oncogenic Ras are primarily determined by the cell of origin and reveal an association between the unlocking of phenotypic plasticity and the onset of tumour-promoting inflammation.

## Main

The Ras gene family comprises the most commonly mutated oncogenes in human cancers^1^. Paradoxically, Ras driver mutations have also been found in normal healthy human tissues^2,3^, indicating that mutations alone are insufficient for tumourigenesis. Non-genetic factors such as intrinsic cellular state and local tissue microenvironment may be instrumental for tumour promotion^4–6^. For example, Ras-driven tumourigenesis can be accelerated by wounding or inflammation^6–8^. Indeed, cancer associated inflammation has long been known to promote tumourigenesis^9,10^. Within established tumours, myeloid cells are reprogrammed to support tumour growth^11,12^, however, little is known about the immediate myeloid response following the first oncogenic mutation. Modelling tumour initiation in zebrafish larvae has enabled *in vivo* visualisation of myeloid responses, showing that oncogenic Ras can trigger cancer-promoting inflammation within hours of oncogene activation^13,14^, although the underlying mechanisms remain elusive.

Phenotypic plasticity is crucial for tumourigenesis as cancer cells must diverge from tissue homeostasis that imposes limits upon cell fate and proliferation^15^. Cells carrying oncogenic mutations initially show increased proliferation but can later undergo differentiation, highlighting differentiation as an early barrier to tumourigenesis^16–18^. Although phenotypic plasticity, such as dedifferentiation or transdifferentiation, is recognised as a requirement for malignant conversion^15,19–21^, most existing studies focus on established tumours or preneoplastic lesions occurring weeks post oncogene activation at which point plastic cell states are already observed. The mechanisms that first unlock phenotypic plasticity and the timing of its acquisition remain to be fully described.

Here we used single cell transcriptomics and *in vivo* live imaging to examine the cellular state of nascent preneoplastic cells (PNCs) and the inflammatory response throughout the first 24 hours following oncogenic HRAS^G12V^ expression in the zebrafish epidermis. Our approach enabled us to determine that basal keratinocytes possess a permissive cellular state for oncogenic Ras driven phenotypic plasticity. Oncogenic Ras induced partial EMT and the acquisition of cancer stem cell features, but only in a subset of basal keratinocytes, demonstrating the importance of non-genetic factors intrinsic to the cell of origin. Cross species data integration revealed a close resemblance between these PNCs and highly invasive cells found at the leading edge of human tumours, showing that malignant potential can emerge within 24 hours of RAS-induced tumour initiation. This subset of PNCs also expressed a unique profile of cytokines implicated in the activation of a systemic neutrophil response. Neutrophils enhanced the proliferation of PNCs, with both immature and mature neutrophils undergoing activation and recruitment to the local PNC microenvironment, similar to the neutrophil response recently described in human tumours.

Our findings identify the intrinsic state of the cell of origin as the primary determinant in driving both phenotypic plasticity and intrinsic tumour-promoting inflammation within the basal epidermis.

## Results

### Cell of origin determines response to oncogenic Ras

HRAS^G12V^ is a common mutation in cutaneous squamous cell carcinoma (cSCC)^22^ and leads to tumour formation in the mouse epidermis^23^. To study Ras-driven tumour initiation we used a tamoxifen-inducible transgenic zebrafish model^24^, wherein the human HRAS^G12V^ oncogene is conditionally expressed within the epidermis under the control of the *krtt1c19e* promoter (Fig. 1A). The zebrafish larval epidermis consists of a superficial periderm and basal keratinocyte layer, the latter of which gives rise to all epidermal lineages within adult zebrafish^25^. The *krtt1c19e* promoter is active within all basal keratinocytes and a small proportion of superficial keratinocytes^25^, therefore, preneoplastic cells within Tg(*krtt1c19e*:KALTA4-ER^T2^; UAS:EGFP-HRAS^G12V^) larvae (hereafter referred to as HRAS larvae) originate from both epidermal layers. Control keratinocytes from Tg(*krtt1c19e*:KALTA4-ER^T2^; UAS:EGFP-CAAX) larvae (hereafter referred to as CAAX larvae) express membrane bound EGFP. The UAS promoter provides mosaic expression of the HRAS^G12V^ or CAAX transgenes throughout the epidermis (Fig. 1B)^26^.

**Figure 1.**
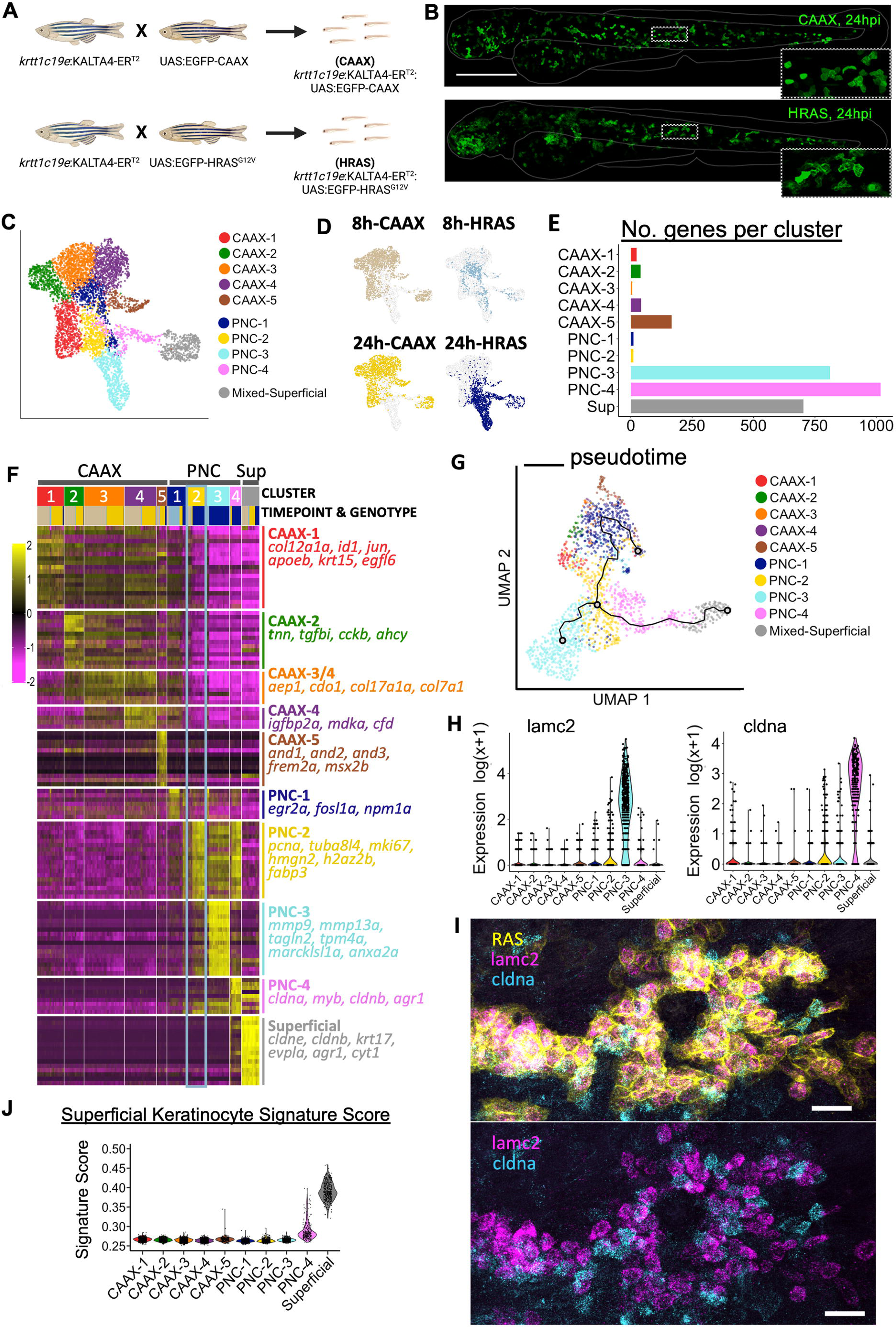
Syngeneic preneoplastic cells diverge to distinct transcriptomic states with 24 hours of HRAS^G12V^ induction. **A,** Schematic showing the breeding of the transgenic HRAS tumour initiation model. HRAS larval genotype (*krtt1c19e:KALTA4-ER^T2^; UAS:EGFP-HRAS^G12V^*), control CAAX larval genotype (*Krtt1c19e:KALTA4-ER^T2^; UAS:EGFP-CAAX*). **B,** Fluorescent images of CAAX and HRAS larvae at 24 hours post induction (hpi) with 4-hydroxytomoxifen. Scale bar= 200µm **C,** UMAP showing scRNA-seq clusters of keratinocytes derived from CAAX larvae (CAAX1 -5) and PNCs derived from HRAS larvae (PNC1-4) at 8 and 24hpi. **D**, UMAP visualisation of keratinocytes by genotype and timepoint. **E,** The number of differentially upregulated genes versus all other clusters, indicates the PNC-3 and PNC-4 clusters are highly distinct populations (pairwise Wilcoxon rank sum tests, genes filtered by detection in >10% of cells within at least one cluster per pair, FDR < 0.05). **F,** Heatmap showing the scaled and centred expression of the top 30 enriched genes per cluster (Welch’s T test between each cluster versus sample. Genes were filtered by detection in >10% of cells within cluster, FDR < 0.05, fold-change >1, and ordered by fold change). **G,** Pseudotime analysis performed on PNCs from 8 and 24 hpi. Black line represents pseudotime trajectory originating from 8 hpi (PNC-1) and reaching a branchpoint within PNC-2. **H,** Violin plots showing the normalised expression of selected markers for PNC-3 (*lamc2*) and PNC-4 (*cldna*). **I**, Whole-mount HCR fluorescence in situ confirms that PNC cluster markers, *lamc2* (magenta) and *cldna* (turquoise), are expressed by distinct populations of PNCs (yellow), within HRAS larvae at 24hpi. Scale bar= 30µm. Lower panel shows that the expression of of *lamc2* and *cldna* is mutually exclusive. **J,** Cells within each cluster were scored for their expression of a “superficial keratinocyte” gene signature, indicating the upregulation of superficial keratinocytes gene within the PNC-4 cluster.

Previously, we found that HRAS^G12V^ expression within the larval epidermis triggers hyperproliferation, morphological changes and an inflammatory response within 24 hours post induction (hpi)^24,27^. Therefore, we isolated epidermal cells and myeloid cells from the same animal by FACS at 8 and 24 hpi to perform single cell RNA sequencing (scRNA-seq), aim to capture the immediate cellular response to oncogenic Ras and host inflammatory cell response to RAS driven preneoplastic initiation (Extended Data Fig. 1A). Cell types were identified by unsupervised clustering and expression of known markers (Extended Data Fig. 1B-D). We found three clusters of basal keratinocytes, one of which was composed almost exclusively of cells derived from HRAS larvae, indicating the emergence of a novel Ras-driven cell state (Extended Data Fig. 1E-F). In contrast, we found a single cluster of superficial keratinocytes composed equally of cells from both CAAX and HRAS larvae. This suggests that basal keratinocytes are more susceptible to Ras-driven changes whilst superficial keratinocytes possess a more fixed cellular identity. Furthermore, differential expression analysis showed >2,000 genes were differentially expressed in basal keratinocytes following HRAS^G12V^ expression compared to only 462 differentially expressed (DE) genes in superficial keratinocytes (Extended Data Fig. 1G).

### Syngeneic preneoplastic cells diverge towards distinct cellular states within 24 hours of oncogene activation

To examine keratinocytes and PNCs in detail we sub-clustered these cells (Fig. 1C). This resulted in a clear separation of basal-derived PNCs from control cells, whereas both CAAX and HRAS superficial keratinocytes remained clustered together (Fig.1 D). Control basal keratinocytes (clusters CAAX-1 to -5) maintained stable cellular states over time (Fig. 1D). In contrast, PNCs showed substantial transcriptomic changes compared to CAAX keratinocytes by 8hpi (PNC-1) and continued to change further over time (PNC-2 to -4) (Fig. 1D). Pairwise differential expression showed that PNC-1 and PNC-2 had very few specific DE genes, indicating a continuous transition (Fig. 1E). In contrast, PNC-3 and PNC-4 were highly distinct with 811 and 1,017 specific DE genes respectively (Fig. 1E).

Permissive DE analysis (see methods) revealed that all basal-derived PNC clusters upregulated cell cycle-related genes, such as *pcna*, *tubb2b* and *mki67,* consistent with the hyperproliferation of PNCs previously described^24^ (Fig. 1F; Supplementary Table 1). Additionally, all basal-derived PNC clusters downregulated control CAAX cluster-specific genes, suggesting a departure from normal development. Immediate Ras/Erk target genes, such as *egr2a* and *fosl1a*^28^ (Fig. 1F), were enriched in PNC-1 capturing the early response to Ras induction at 8hpi. PNC-3 enriched genes included *mmp9* and *mmp13a*, and actin binding proteins associated with cancer cell migration, such as *tpm4*^29^, *tagln2*^30^ and *anxa2*^31^. In contrast, PNC-4 enriched genes included *myb*, a key regulator of terminal differentiation in the epidermis^32^ and *cldna*, a marker of intermediate keratinocytes^33^, suggesting that these cells undergo differentiation to suprabasal fate. In support of this, there was also a noteworthy overlap between genes enriched within PNC-3 and the superficial keratinocyte cluster, including superficial markers, such as *cyt1*, *cytl1*, *cldne*, *cldnb* and *krt17* ^33,34^. Finally, whilst PNC-2 lacked specifically enriched genes, this cluster expressed low levels of both PNC-3 and PNC-4 enriched genes (Fig. 1F), suggestive of an intermediate state. Indeed, pseudotime analysis identified a branch point within PNC-2 at which PNCs diverge towards the two distinct cell states (Fig. 1G).

To determine whether distinct subpopulations of PNCs were present *in vivo*, we used HCR^TM^ fluorescence in situ hybrisation (HCR FISH) to visualise gene expression. We confirmed that the expression of PNC-3 marker *lamc2* and PNC-4 marker *cldna* were mutually exclusive (Fig. 1H, I). Moreover, within the same PNC clone, some *cldna* positive PNCs were positioned apically to *lamc2* positive PNCs (Supplementary Video 1). This is consistent with the enrichment of intermediate and superficial keratinocyte markers within PNC-4 (Fig. 1J) and confirms their differentiation to suprabasal fate^35^.

These findings show that syngeneic PNCs, even when derived from the same cell type, can undergo divergent cell state transitions. Whilst some remained aligned with normal tissue differentiation occurring during homeostasis (PNC-4), others unlocked unique cellular states distinct from normal epidermis (PNC-2 / PNC-3).

### Subsets of Ras-driven PNCs undergo dedifferentiation to an embryonic-like state

To explore transcriptomic differences between PNCs and normal epidermis, we performed pairwise pseudo-bulk comparisons between each of the 24hpi PNC clusters versus control basal keratinocytes (Supplementary Table 2). The 100 most downregulated genes in all three clusters (PNC-2, 3 and 4) included numerous basal keratinocyte markers, collagens and keratins (Fig. 2A). This downregulation is expected in PNC-4 cells due to their differentiation process. However, the absence of differentiation markers in PNC-2 and PNC-3, despite similar downregulation, suggests a potential dedifferentiation occurring within these clusters.

**Figure 2.**
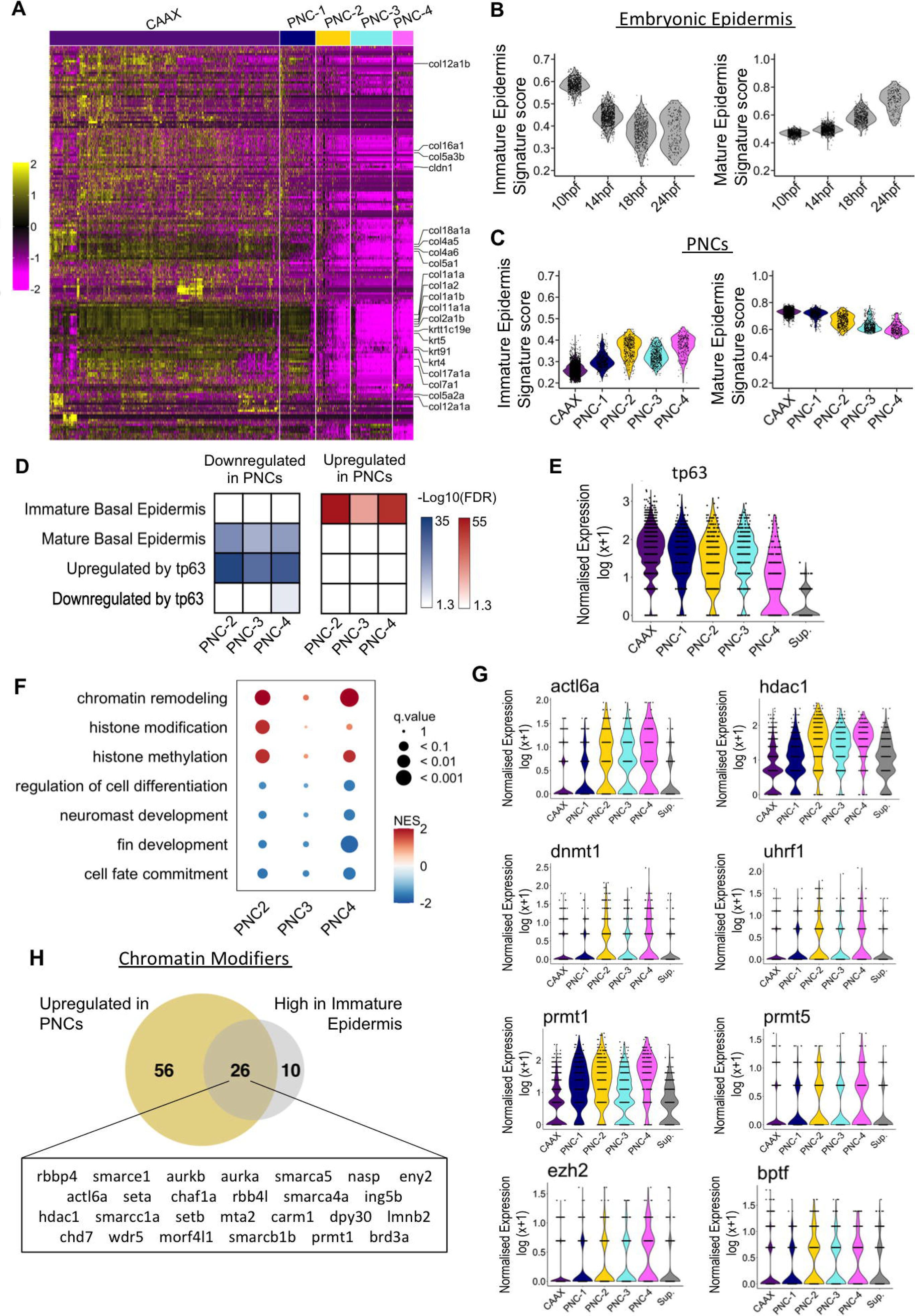
HRAS^G12V^ expression causes rapid dedifferentiation to an embryonic-like state, accompanied by the upregulation of epigenetic modifiers. **A,** Heatmap depicting the scaled and centred expression of the top 100 downregulated genes in each PNC cluster, indicates a downregulation of characteristic basal keratinocyte genes (PNC clusters were compared with all control basal keratinocytes by pseudobulk differential expression analysis. Genes were filtered by detection in >10% of cells within at least one group, FDR < 0.05, fold-change > |1.5|). **B,** scRNA-seq data from zebrafish embryonic development (Wagner et al. 2018) ^39^ was reanalysed to derive gene signatures relating to immature epidermis versus mature epidermis. The violin plots depict signature scores within epidermal cells from the original data sets at successive timepoints during embryonic development. **C,** The gene signatures described in B were used to score PNCs, indicating downregulation of mature epidermal genes and re-expression of immature epidermal genes. **D,** Over-representation analysis of the immature and mature epidermal signatures depicts the significance of their enrichment within genes downregulated (blue) and upregulated (red) within PNC clusters at 24hpi. **E,** Violin plots show normalised expression of *tp63* in control (CAAX) basal keratinocytes, PNC clusters and superficial keratinocytes (Sup.) **F,** Gene Set Enrichment Analysis of Gene Ontology “Biological Processes” for each PNC cluster vs. control basal keratinocytes indicates upregulation of epigenetic modifiers. Colours represent normalised enrichment score, radius represents q-value. **G,** Violin plots showing the normalised expression of histone modifiers that are both upregulated within PNCs and known to contribute to the maintenance of stemness within the epidermis. **H,** Venn diagrams depicts the overlap between the chromatin modifiers upregulated within PNCs and those within the immature epidermis gene signature.

In many aggressive cancers, cells travel backwards along the Waddington landscape and begin to re-express gene programmes reminiscent of embryonic precursors^20,36–38^. To determine whether this occurs within Ras-driven PNCs, we derived gene sets from zebrafish embryonic development^39,40^ (Fig. 2B). Immature epidermal genes were upregulated in all PNC clusters, whilst mature epidermal genes were downregulated (Fig. 2 C&D). To investigate this further we inspected the expression of *tp63* target genes, a master regulator of the epidermal lineage^41^. Genes known to be positively regulated by *tp63* were downregulated within PNCs (Fig. 2D) despite sustained *tp63* expression (Fig. 2E). *Tp63* also acts to repress neuroectoderm genes during embryogenesis^40^, therefore, we questioned whether dedifferentiation might open a gateway for transdifferentiation. However, genes known to be negatively regulated by *tp63*, including neuroectoderm genes, were not differentially expressed within PNCs (Fig. 2D). This indicates dedifferentiation towards an embryonic-like epidermal state without transdifferentiation to neuroectoderm fate.

Furthermore, Gene Set Enrichment Analysis showed downregulation of genes related to differentiation, development and cell fate commitment, whilst gene sets related to epigenetic programming were upregulated (Fig. 2F). Upregulated epigenetic regulators included *dnmt1*, *uhrf*, *hdac1*, *ezh2*, *actl6a*, *prmt1*, *prmt5* and *bptf*^42–44^ (Fig. 2G), which are known to suppress differentiation and maintain basal stem cell self-renewal. Many of these chromatin modifiers were also part of the immature epidermis signature (Fig. 2H). Thus, oncogenic Ras expression led to dedifferentiation within a subset of basal keratinocytes, likely regulated at the epigenetic level.

### A subpopulation of PNCs undergoes partial EMT and resembles cancer stem cells

Epithelial-to-Mesenchymal transition occurs in tumours and is linked to invasion and metastasis^45^. The genes specifically upregulated within the PNC-3 cluster are strongly indicative of EMT. These include genes related to cell migration, such as the arp2/3 complex required for lamellipodia formation, myosins involved in rear contractility, components of migratory signalling pathways and key integrins (Fig. 3A, Extended Data Fig. 2A). Cell surface receptors related to EMT and invasion in cancer such as *cd276*^46^, *cd81a*^47^, *nrp1a/b*^48^ were also upregulated. Upregulation of EMT genes was validated by qPCR of FACS sorted PNCs (Extended Data Fig. 2B). However, the PNC-3 cluster retained epithelial markers, *cdh1* (E-cadherin), *epcam*, and *tjp1a* (zonal occludins 1) and lacked some mesenchymal markers such as *vim* (vimentin) and *cdh2* (N-cadherin) (Fig. 3A & Extended Data Fig. 2C), indicating only partial EMT (pEMT). Consistent with this finding, the only EMT transcription factor upregulated in PNC-3, *snai2* (Extended Data Fig. 2C), was previously found to drive pEMT in cancer^49,50^.

**Figure 3.**
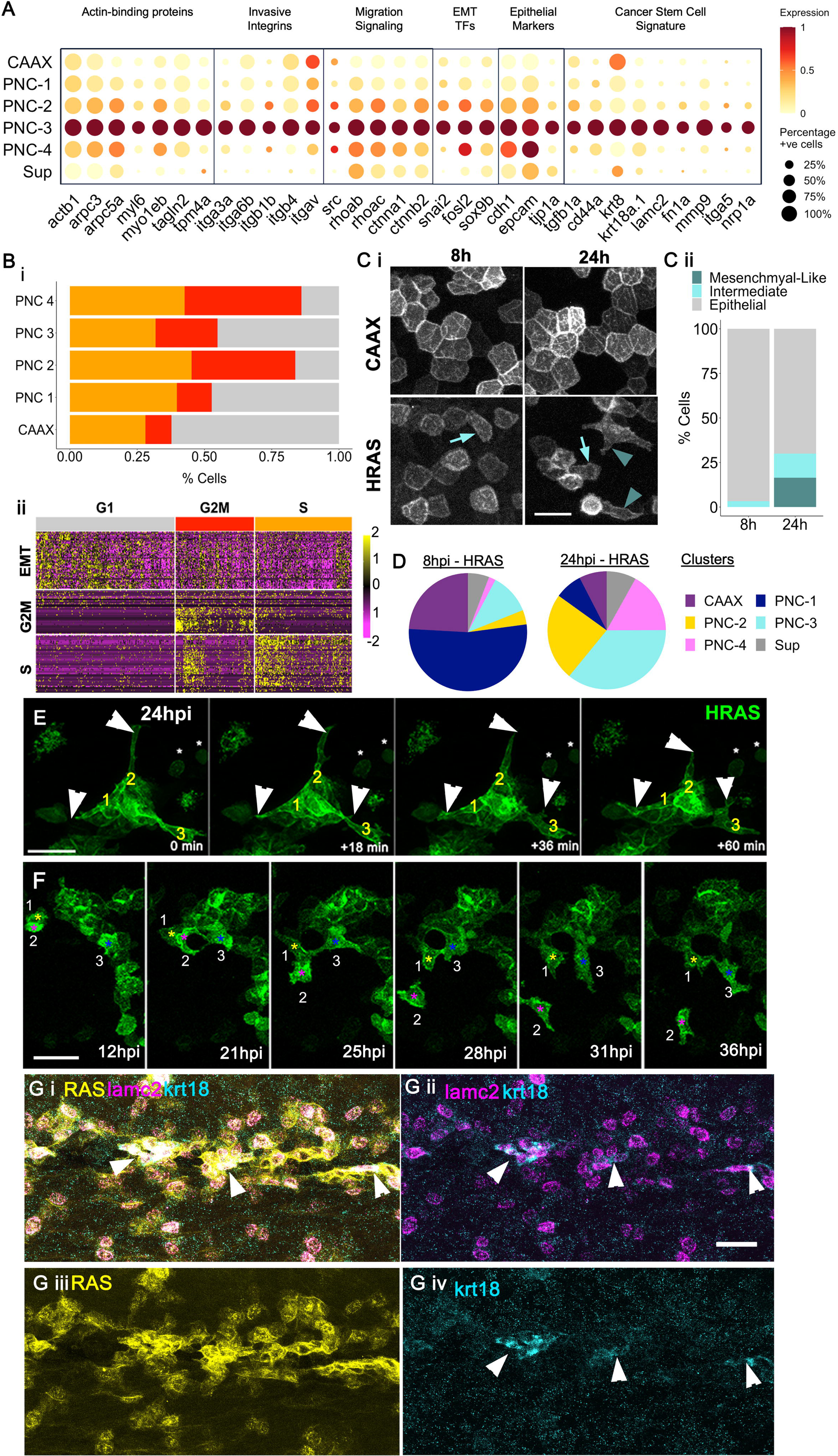
PNC-3 undergoes partial Epithelial to Mesenchymal transition (pEMT), acquired ability to migrate and harbours cells positive for a cancer stem cell marker. **A,** Dot plot showing the scaled expression of genes relating to EMT and cancer stem cells, selected by significant upregulation in PNC-3 (Genes were filtered by detection in >10% of cells within PNC-3, FDR < 0.05, fold-change > 1.25). **B**, **(i)** Cell cycle scoring algorithms were used to infer the cell cycle phase from the scRNA-seq data, showing that the PNC-3 cluster is less proliferative than other PNC clusters at 24hpi. **(ii)** Heatmap shows the scaled and centred expression of genes relating to EMT and the cell cycle with the cells of the PNC-3 cluster only. Cell are grouped by cell cycle phase, indicating that cells with the highest expression of EMT genes have the lowest expression of cell cycle genes. **C, (i)** Representative confocal live images of control basal keratinocytes (CAAX) and PNCs (HRAS) at 8hpi and 24hpi, show that PNCs undergo morphological changes over time. Arrowheads highlight PNCs with mesenchymal-like features such as elongation and protrusions. Arrows indicate PNC with intermediate morphology. Scale bar= µm. **(ii)** PNCs were manually classified by their morphology as epithelial, intermediate, or mesenchymal-like and quantified as a percentage of total PNCs (n = 10 larvae per condition, av. number of PNCs analysed per larva = 106). **D,** Pie charts depicting the percent of cells belonging to each cluster within the 8hpi and 24hpi HRAS samples. **E,** Individual frames from time-lapse confocal imaging of PNCs from 24-25hpi (Supplementary Movie 2). Arrowheads indicate protrusions that are extended and contracted over time. Asterisks indicate cells that maintain a round shape and lack membrane dynamics. Scale bar= 30µm. **F,** Individual frames from time-lapse confocal imaging of PNCs from 12-36hpi (Supplementary Movie 3) showing that PNCs become motile and start to migrate at approx. 24hpi. Each coloured asterisks indicates a cell that changed position over time. Scale bar= 30µm. **G,** Whole-mount HCR fluorescence in situ showing the expression of cancer stem cell marker keratin 18 (turquoise), within a sub-population of PNC-3 cells (*lamc2*, magenta), arrowheads indicate groups of *krt18* positive cells. Scale bar= 30µm.

Partial EMT has been associated with cancer stem cells (CSCs), with hybrid E/M cells possessing greater plasticity and tumour initiating capacity than cells at the extremes of the E/M spectrum^51^. Therefore, we questioned whether the PNC-3 cluster may contain additional cancer stem cell (CSC) features. Indeed, CSC markers *cd44a* and *thy1*^52^ were specifically upregulated within PNC-3 (Fig. 3A, Extended Data Fig. 2D), alongside markers of a CSC population recently identified in a mouse cSCC model^53^ (Fig. 3A). Akin to normal stem cells, CSCs undergo a low rate of proliferation^53^. Inference of cell cycle phase suggested that, although all PNC clusters had an elevated rate of proliferation in comparison to control cells, PNC-3 was the least proliferative cluster (Fig. 3Bi). In fact, the expression of pEMT gene program was inversely correlated with the expression of cell cycle genes (Fig. 3Bii) and cell cycle inhibitors were upregulated specifically within PNC-3 (Extended Data Fig. 2E).

To ascertain whether evidence of pEMT could be observed *in vivo*, we performed live confocal imaging. In comparison to the regular polygonal shape of control basal keratinocytes, PNCs began to undergo morphological changes by 8hpi, some PNCs losing the polygonal shape entirely by 24hpi (Fig. 3C). PNCs became either rounded or stretched, the latter of which possessed protrusions resembling lamellipodia and were classified as “mesenchymal-like” (Fig. 3C). At 24hpi, live imaging revealed 30% of PNCs were mesenchymal-like (Fig. 3C), similar to the proportion of PNCs belonging to the PNC-3 cluster (39%) (Fig. 3D). All mesenchymal-like PNCs dynamically extended and retracted their protrusions, whilst some dissociated from their neighbouring cells and migrated laterally (Fig. 3E&F, Supplementary Video 2, Supplementary Video 3). These findings challenge current conceptual understanding of tumourigenesis, clearly demonstrating that traits typically associated with late-stage malignant tumours, such as EMT and cell migration, can manifest within just 24 hours of oncogenic Ras activation.

Furthermore, we confirmed the expression of CSC marker *krt18a*^53^ *in vivo*. *Krt18a^+^* PNCs were a small subset of *lamc2^+^* PNCs, typically found adjacent to one another within small groups suggestive of self-renewing clonal expansion (Fig. 3G). Thus, we have identified a subset of PNCs resembling cancer stem cells, further showing that not all PNCs are created equal, and some may have greater plasticity and malignant potential from their inception.

### Distinct PNC states correspond to subpopulations found with human tumours

Having identified distinct cellular states, including cancer stem cell like features, amongst preneoplastic cells, we next sought to determine whether such states were conserved in human disease. To address this, we performed a comparative analysis with a scRNA-seq dataset from human cSCC patients^54^ (Accession number: GSE144240). We used LIGER^55^ to directly integrate our 24hpi PNCs & matched CAAX cells with basal cells from 10 SCC patient tumours and matched healthy tissue. The integrated data was jointly clustered to identify shared cellular states.

Control basal keratinocytes from zebrafish co-clustered with human basal keratinocytes from the healthy controls, indicating successful data integration (Fig. 4A&B). PNC-4 cells co-clustered with differentiated tumour cells, whilst PNC-3 cells co-clustered with an invasive “tumour specific” population found at the leading edge of tumours (Fig. 4&B). Akin to PNC-3, the “tumour specific” cells lacked basal keratinocyte markers, upregulated *snai2,* underwent EMT, and had a low proliferation rate (Fig. 4A&B).

**Figure 4.**
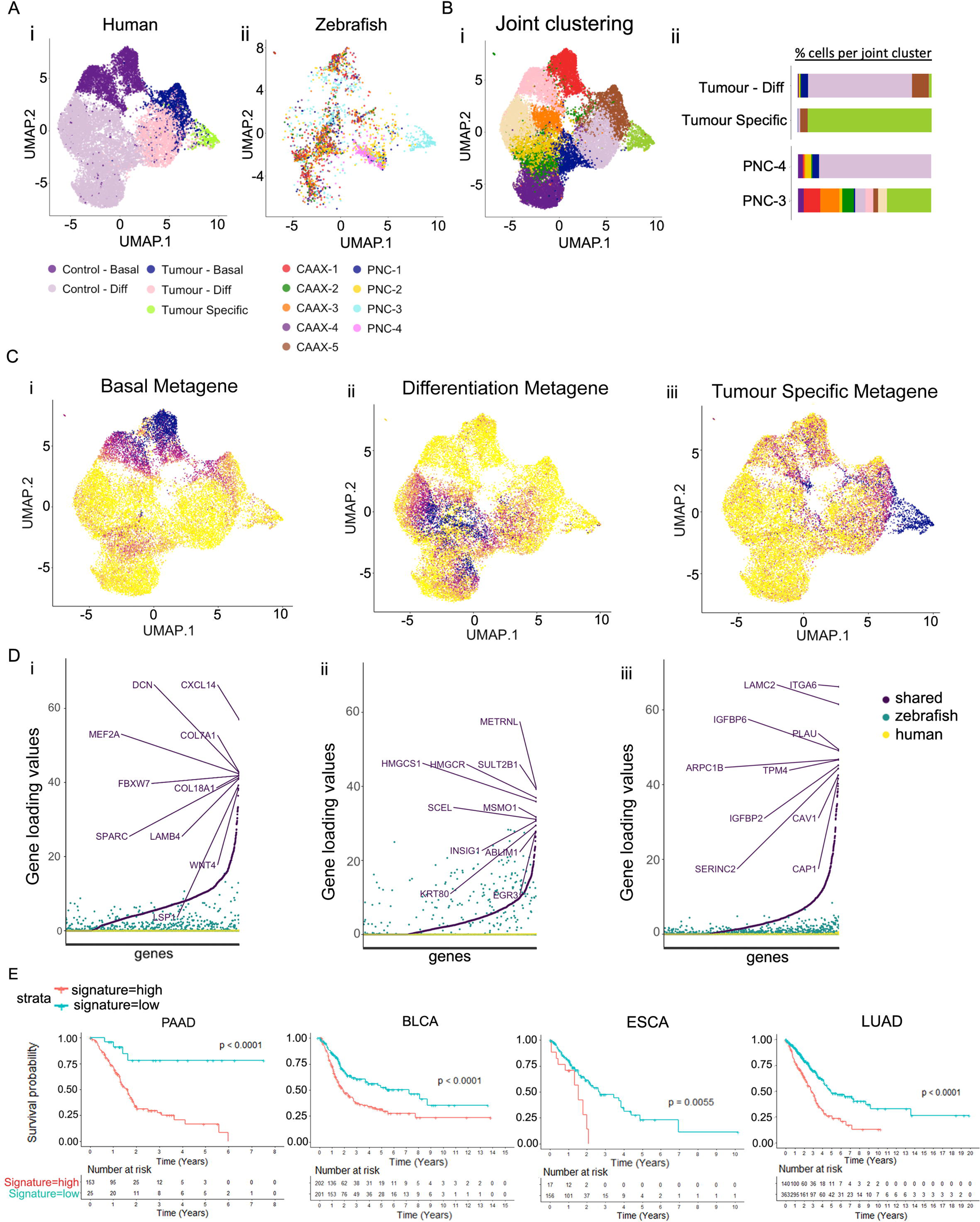
The pEMT subpopulation of PNCs closely corresponds to the invasive cells found at the leading edge of human tumours and is associated with reduced patient survival. **A,** CAAX keratinocytes and PNCs were integrated with a scRNA-seq dataset of cutaneous squamous cell carcinoma and matched healthy skin (Ji et al. 2020). UMAPs were derived following integration, colours depict (i) the original clusters of human cSCC cells and healthy skin from Ji et al, (ii) the original clusters of zebrafish PNCs and basal keratinocytes first depicted in Figure 1C. **B,** The integrated datasets were jointly clustered, (i) UMAP depicts new clusters following integration, (ii) percentage of cells per joint cluster, indicating that PNC-4 cells cluster together with differentiated cells from human cSCC, whilst PNC-3 cells correspond with a highly malignant tumour specific human cSCC cells. **C,** LIGER analysis identified metagenes shared between human and zebrafish datasets. UMAPs depict the expression of the following metagenes: (i) basal, (ii) differentiation, (ii) tumour specific. **D,** Plots depict the contribution of zebrafish-specific, human-specific and shared genes to the following metagenes: (i) basal, (ii) differentiation, (ii) tumour specific. This indicates that shared genes were the major contributors to these metagenes and the tumour specific metagenes was most highly conserved between species. The top ten contributing genes for each metagenes are annotated. **E,** Kaplan-Meier curves show overall survival in patients across the primary RAS-driven cancer types. Patients are split according to expression of PNC-3 cluster genes (red = high expression, blue = low expression) with patients expressing high levels of the PNC-3 signature exhibiting a significantly reduced overall survival than those with low expression. P-values and hazard ratios were calculated using log-rank tests and Cox proportional hazard models, respectively.

LIGER identified sets of co-regulated genes (metagenes) shared between the zebrafish and human datasets (Fig. 4C&D). These included a “basal” metagene featuring collagens and laminins (Fig. 4Di); a “differentiation” metagene featuring suprabasal markers SULT2B1, KRT80 and EGR3 (Fig. 4Dii); and a “tumour specific” metagene featuring the PNC-3 marker LAMC2 and cell migration genes ITGA6 and ARPC1B (Fig. 4Diii). For each of these metagenes, the contribution from shared genes was greater than that of zebrafish-specific or human-specific genes (Fig. 4D). This was especially true for the “tumour-specific” metagene, indicating that PNC-3 cells are closely related to malignant cells in human tumours.

If PNC-3 cells represent a malignant cellular state that can persist in established tumours, we would expect the markers of this cluster to be associated with poor outcomes in cancer patients. To test this hypothesis, we used PNC-3 marker gene signature (Supplementary table 3) to perform survival analysis using patient data from the Cancer Genome Atlas (TCGA) PanCancer dataset. We found that the PNC-3 signature was associated with poor patient survival in 17 out of the 25 cancer types, particularly those driven by RAS mutations (Fig. 4E) (Hazard Ratios: PAAD, 6.28; BLCA, 1.90; ESCA, 2.57; LUAD, 2.15).

### Neutrophils are the first immune cells to respond following Ras-driven tumour initiation

We and others have shown that HRAS^G12V^ driven PNCs induce an inflammatory response, but the mechanisms remain unclear^13,56^. Our scRNA-seq data revealed upregulation of cytokines and chemokines within PNCs in as early as 8hpi (PNC-1): *il1b*, *csf3b*, *il34*, *cxcl8b.1*, *cxcl18a.1*, *il11b*, and *mif* (Fig. 5A). This was sustained at 24hpi (Fig. 5A) and upregulation of *il1b, cxcl8* and *cxcl18* was confirmed by qPCR (Fig. 5B). Additional cytokines were upregulated in PNC -3 and -4 clusters at 24hpi (Fig. 5A). PNC-3 exhibited a unique secretome, co-expressing pro-inflammatory cytokines alongside *tgfb1a* and *inhbaa*/*inhbb*, which have been shown to programme immunosuppressive myeloid cells within tumours^57,58^ (Fig. 5A).

**Figure 5.**
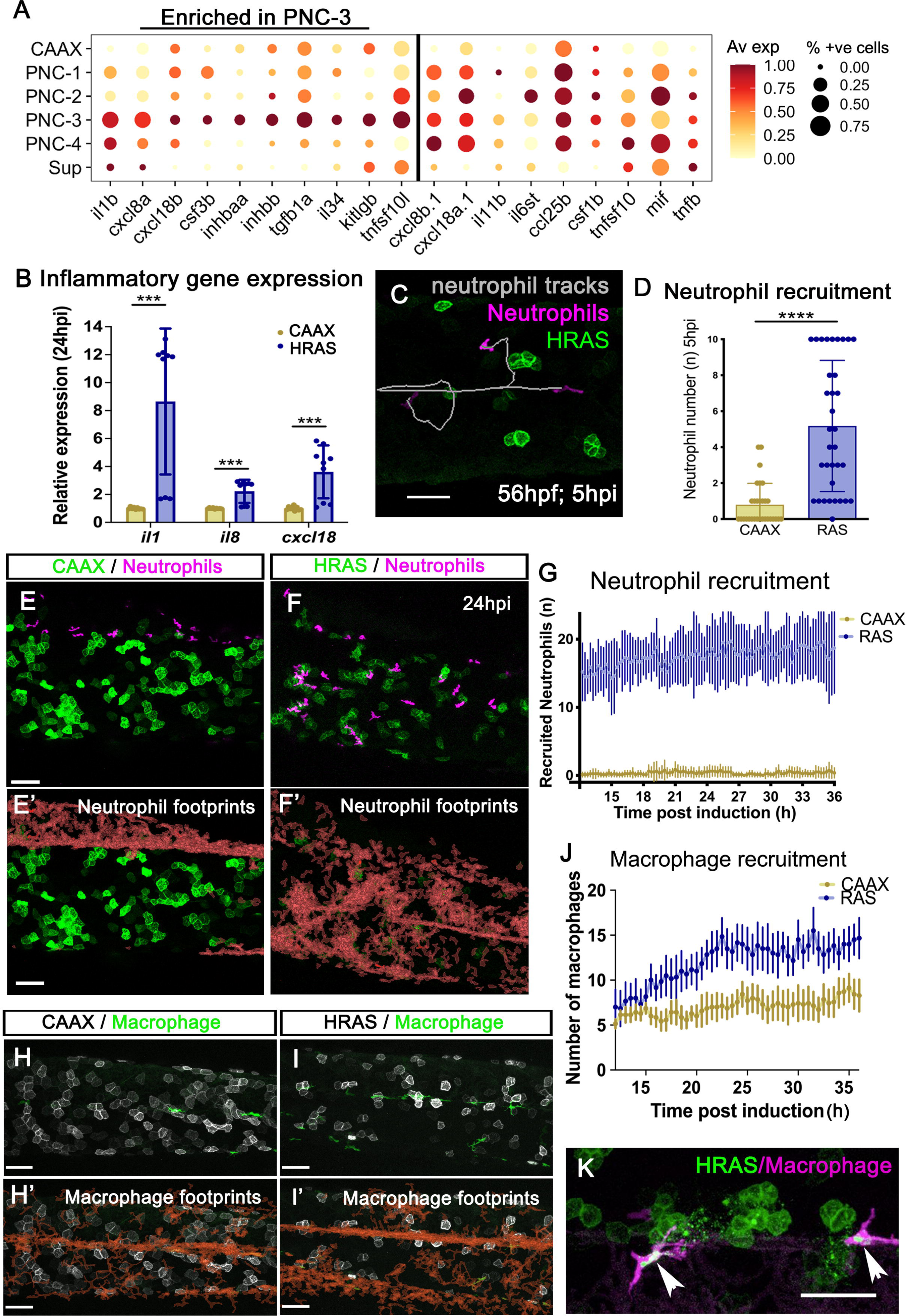
PNCs up-regulate inflammatory cytokines and chemokines, and rapidly recruit both neutrophils and macrophages. **A,** Dot plot show scaled expression of all cytokines and chemokines upregulated in PNCs versus control CAAX keratinocytes, many of which were enriched within the PNC-3 cluster. **B,** qPCR shows the upregulation of pro-inflammatory cytokines and chemokines (*il1*, *cxcl8* and *cxcl18*) in PNCs extracted from HRAS larvae versus control keratinocytes extracted from CAAX larvae (unpaired t test, p<=0.0008, n=9). **C,** Confocal imaging of neutrophils *in vivo* shows recruitment as early as 5hpi. Neutrophil tracks are shown in white (Supplementary Movie 4). Scale bar = 50µm. **D,** Quantification of the number of neutrophils observed within the epidermis of control versus PNC-bearing larvae at 5hpi (Mann-Whitney test, P<0.0001, n>=30). **E,** Confocal image of neutrophils (magenta) and keratinocytes (green) within the epidermis of CAAX larvae at 24hpi (derived from Supplementary Movie 5). **E’**, Red areas indicate neutrophil footprints from 12 to 36 hpi. **F,** Confocal image of neutrophils (magenta) and PNCs (green) within the epidermis of HRAS larvae at 24hpi (derived from Supplementary Movie 6). **F’**, Red areas indicate neutrophil footprints from 12 to 36 hpi. Scale bar= 50µm in E-F’.**G,** Quantification of neutrophil infiltration into the epidermis of control (CAAX) versus PNC-bearing (HRAS) larvae from 12-36 hpi (two-way ANOVA, p<0.0001, n=10). **H,** Confocal image of macrophages (green) and keratinocytes (white) within the epidermis of CAAX larvae at 24hpi (derived from Supplementary Movie 7). **H’,** Footprints (orange) showing macrophage patrolling in the larval epidermis from 12-36 hpi. **I,** Confocal image of macrophages (green) and PNCs (white) within the epidermis of HRAS larvae at 24hpi (derived from Supplementary Movie 8). **I’,** Footprints (orange) show macrophage recruitment to PNCs. Scale bar = 50µm in H-I’. **J,** Quantification of macrophage numbers in the epidermis of control (CAAX) versus PNC-bearing (HRAS) larvae from 12-36hpi (two-way ANOVA, p<0.0265, n>= 6). **K,** A still image from a confocal time-laps (Supplementary Movie 9) depicting macrophages (magenta) engulfing dying PNCs (green), arrowheads highlight PNC material within macrophages. Scale bar = 30µm.

The chemokines upregulated by PNCs included known neutrophil (*cxcl8*, *cxcl18*) and macrophage (*il34*, *csf1b*) chemoattractants (Fig. 5A). Live imaging revealed that neutrophils were recruited to the epidermis of HRAS larvae by 5hpi, the earliest time that PNCs can be detected (Fig. 5C&D, Supplementary Video 4). Neutrophils underwent dynamic interactions with PNCs and continued to accumulate over time (Fig. 5E-G, Supplementary Video 5, 6). In contrast, although macrophages constitutively patrol the healthy epidermis, they did not increase in number compared to that in control fish or interact preferentially with PNCs until >16hpi (Fig. 5H-J,S upplementary Video 7, 8). By 24hpi macrophages could be observed engulfing the debris of dying PNCs (Fig. 5J&K, Supplementary Video 9).

### Macrophages do not resemble pro-tumour TAMs at 24hpi

Within tumours, macrophages form specialised populations that support tumour growth and often have immunosuppressive properties^11^. To assess their response following tumour initiation, we analysed macrophages from our scRNA-seq dataset (Extended Data Fig. 1A-D). Surprisingly, macrophages underwent minimal transcriptomic changes within 24 hours of PNC induction (Extended Data Fig. 3C&E). They mildly upregulated pro-inflammatory genes but did not upregulate alternative activation markers frequently expressed by tumour associated macrophages (TAMs) (Extended Data Fig. 3F). Moreover, macrophage depletion did not affect PNC proliferation at 24 hpi (Extended Data Fig. 3G&H). Thus, macrophages do not play a significant role following preneoplastic initiation in our model within this time frame.

### Ras-driven PNCs elicit systemic changes to granulopoiesis

To assess the neutrophil response following PNC induction we re-clustered neutrophils from the scRNA-seq data. Neutrophils derived from whole CAAX and HRAS larvae separated almost completely at both 8 and 24hpi (Fig. 6A). Since neutrophils were derived from whole larvae, this indicates a rapid systemic response. Control neutrophil clusters were defined by developmental stage, from immature (C1) to mature (C4) (Fig. 6B, Supplementary Table 4). Neutrophils from HRAS larvae similarly clustered according to their maturation, from H1 to H4 (Fig. 6B & Extended Data Fig. 4A&B).

**Figure 6.**
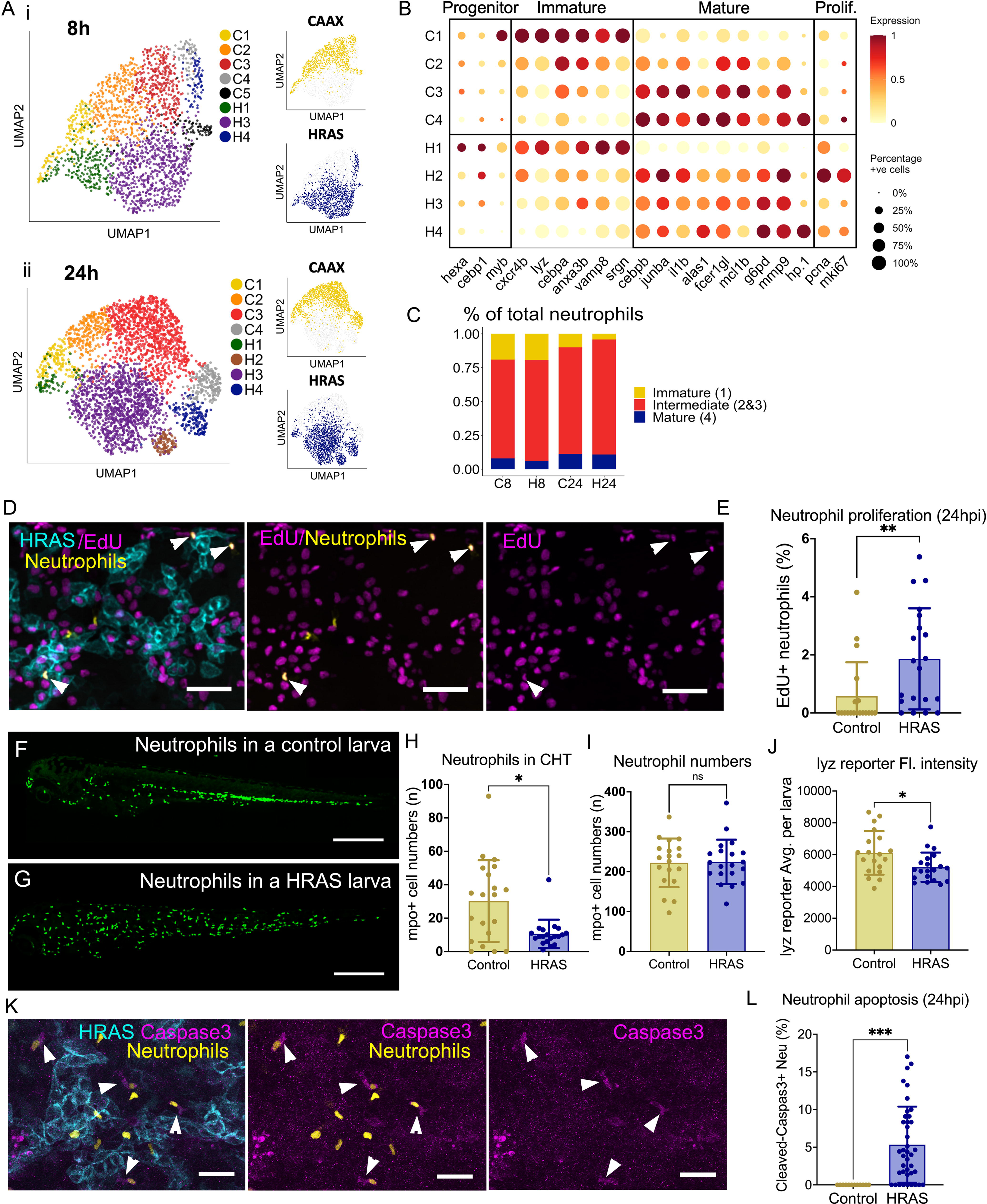
PNCs elicit a systemic neutrophil response resulting in accelerated granulopoiesis. **A,** UMAPs depict the clustering of neutrophils derived from control CAAX larvae and PNC-bearing larvae at (i) 8hpi, and (ii) 24 hpi. **B,** Dot plot representing the expression of genes associated with Granulocytic Myeloid Progenitors, immature neutrophils and mature neutrophils. **C,** Neutrophils within the scRNA-seq data were grouped by their maturity and quantified as a percentage of total neutrophils per samples. **D,** Representative fluorescent confocal images showing HRAS larval epidermis following EdU staining, depicting PNCs (turquoise) neutrophils (yellow) and EdU^+ve^ nuclei (magenta). White arrowhead indicating EdU^+ve^ neutrophils. Scale bar = 30µm. Percentage of EdU^+ve^ neutrophils quantified in E. **E**, Graph showing comparison of percentage of EdU^+ve^ neutrophils between siblings and HRAS larvae (Mann-Whitney test, N>=19, **p=0.0017). **F,** Representative fluorescent confocal image showing mpo^+ve^ neutrophil (green) distribution in a whole wild-type larva. Scale bar=200µm. **G,** Representative fluorescent confocal image showing mpo^+ve^ neutrophil (green) distribution in a whole HRAS larva. Scale bar=200µm. **H,** Graph showing comparison of neutrophils reside in the CHT area between wild type siblings and HRAS larvae. (Mann-Whitney test, N=20, *p=0.0118.) **I,** Graph showing comparison of total neutrophils between siblings and HRAS larvae (unpaired t test, N>=19, p=0.888.). **J** Graph showing comparison of average fluorescent intensity of neutrophil population between siblings and HRAS larvae. (unpaired t test, N>=19, *p=0.0203). **K,** Confocal fluorescent images showing anti-cleaved-caspase staining of HRAS larval epidermis, PNCs (turquoise), neutrophils (yellow) and cleaved-caspase3 (magenta). White arrowheads indicate caspase^+ve^ neutrophils. Scale bar = 30µm. Percentage of cleaved-caspase 3 +ve neutrophils quantified in L. **L,** Graph showing comparison of apoptotic neutrophils between siblings and HRAS larvae (Mann-Whitney test, N>= 10, ****p<0.0001).

Although neutrophils from HRAS larvae spanned the full spectrum of maturation seen in control larvae, many genes related to neutrophil development were differentially expressed in H1-H4 clusters compared to their control counterparts at 24hpi (Fig. 6B). The transcription factor that governs steady state granulopoiesis, *cebpa*^59^, was downregulated in all HRAS neutrophil clusters (Fig. 6B), whilst the master regulator for emergency granulopoiesis, *cebpb*^60,61^, was highly upregulated in the H2 cluster alongside proliferation markers, *pcna* and *mki67* (Fig. 6B). Immature neutrophil genes, *lyz*, *cxcr4b*^62^ and *myb* ^63,64^ were downregulated within the H1 progenitor clusters. Moreover, the proportion of immature neutrophils (C1 & H1) was decreased in HRAS vs. CAAX samples at 24hpi, while mature neutrophil levels remained constant (Fig. 6C). This indicates that PNC induction triggered an acceleration of early granulopoiesis and expansion of intermediate stage neutrophils (H2 & H3). Indeed, the PNC cytokine expression profile described above included upregulation of *csf3b* (G-CSF), a known trigger of emergency granulopoiesis^60^ (Fig. 5A)

Using EdU incorporation assay, we detected increased EdU^+^ lyz^+^ neutrophils in PNC-bearing larvae (Fig. 6D&E), confirming accelerated granulopoiesis following PNC initiation. Imaging neutrophil distribution in the whole larvae, we found that the majority of neutrophils within HRAS larvae emigrated from the Caudal Hematopoietic Tissue (CHT) and entered the epidermis (Fig. 6F-H). However, the total number of neutrophils remained unchanged (Fig. 6I), indicating that increased granulopoiesis might be balanced by neutrophil loss. Cleaved-caspase 3 staining confirmed that some neutrophils recruited to the epidermis of HRAS larvae underwent apoptotic cell death (Fig. 6K&L). This is reminiscent of emergency granulopoiesis, during which an initial loss of neutrophils is compensated by accelerated neutrophil production^60^. Furthermore, the mean fluorescence intensity from whole body neutrophil imaging using the *lyz* reporter was significantly decreased in PNC-bearing larvae, validating the decreased *lyz* expression observed in the scRNA-seq data (Fig. 6J) and again indicating the accelerated development of neutrophils.

### Both immature and mature neutrophils are recruited to PNCs and respond in a similar manner

In addition to accelerated granulopoiesis, *cs3fb* (G-CSF), expressed by the PNCs, can cause premature release of immature neutrophils from haematopoietic tissues via the downregulation of Cxcr4^65,66^. Indeed, *cxcr4b* was downregulated within immature neutrophils from PNC-bearing larvae (Fig. 6B). To directly visualise immature versus mature neutrophils *in vivo*, we used *lyz:lifAct-*mTurquoise2a and *myd88*:DsRed reporters^67^ (Fig. 7A). As previously described, *lyz* is most highly expressed within immature neutrophils and downregulated as neutrophils mature^64^. *Myd88* expression increases as neutrophils mature and we confirmed *myd88* reporter expression negatively correlated with *lyz* reporter expression in PNC-bearing larvae (Fig. 7A-C). Live confocal imaging showed that both mature neutrophils (*myd88*^hi^ / *lyz^l^*°) and immature neutrophils (*myd88*^lo^ / *lyz*^hi^) were recruited to PNCs but immature neutrophils had a lower speed of migration and longer retention time near PNCs in comparison to mature neutrophils (Fig 7. D&E, Supplementary Movie 10), indicating behavioural differences between these populations.

**Figure 7.**
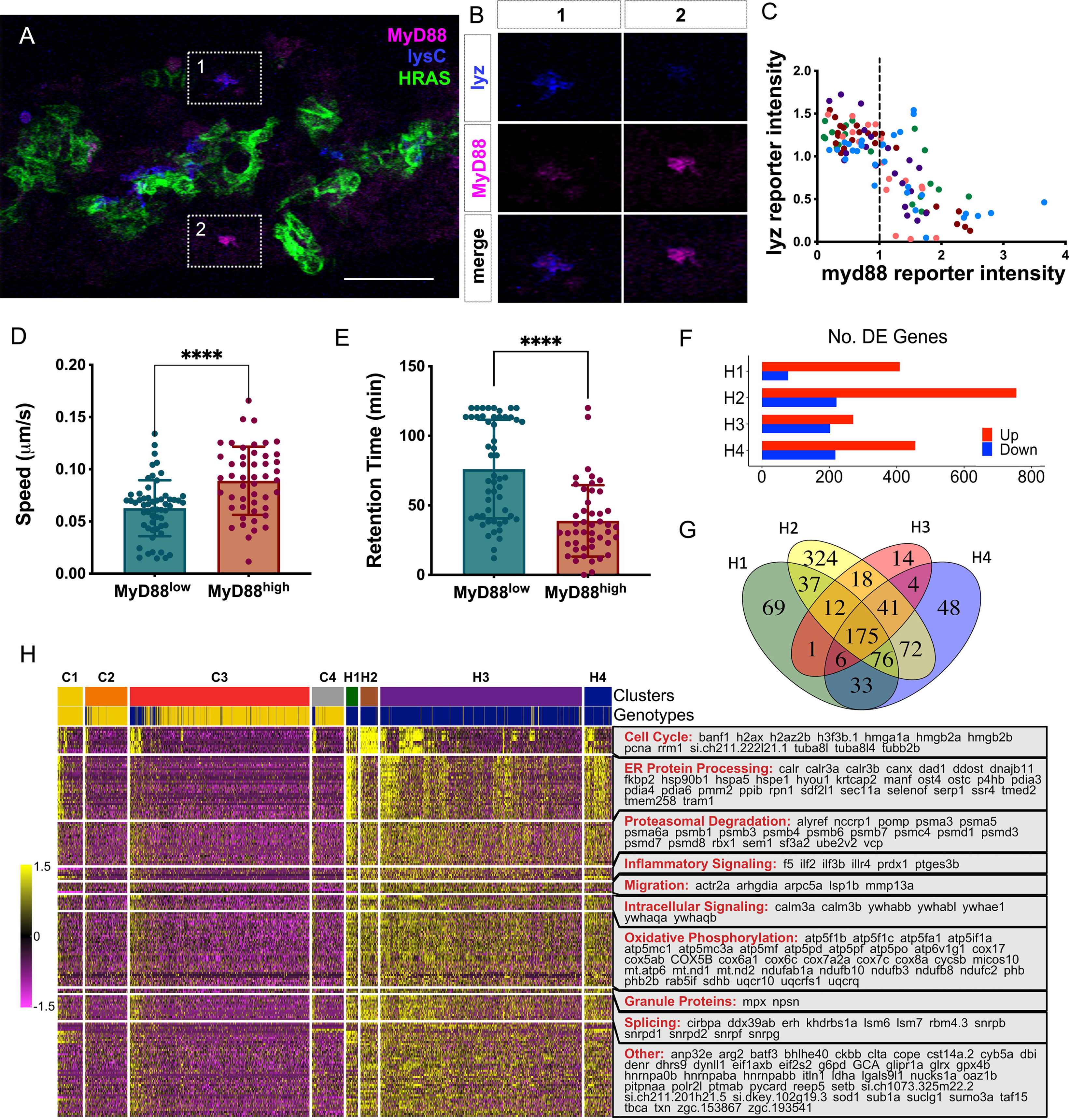
Both immature and mature neutrophils respond to PNC induction but remain distinctive behaviour. **A,** Still image of a time lapse movie showing HRAS larva epidermis, PNCs (green), recruited neutrophils displaying different levels of *lyz*:lifAct-mturquoise2a reporter (blue) intensity; and *myd88*:dsred2 (magenta) intensity. Scale bar = 100µm. **B,** Zoom in single channel images from box1, 2 in A, showing presence of both 1, immature neutrophils (*myd88*^lo^ / *lyz*^hi^) and 2, mature neutrophils (*myd88*^hi^ / *lyz^l^*°). **C,** Scatter plot of individual neutrophils recruited within a 2-hour time lapse movie at 24hpi, showing quantification of mean fluorescent intensity of reporters in neutrophils according to mTurquoise (Y axis) and DsRed (X axis) channels. Each colour represents neutrophils from an individual larva. To account for variation of the same reporter between different larva, due to differences in transgene copy number, mean intensity values of each neutrophil were normalized to average intensity of all neutrophils within each larva. Consequently, plotted values represent relative intensity where X/Y=1 represents the neutrophil average intensity within each larva. Dashed line at X=1 in represents the value separating the two neutrophil subsets compared in the subsequent analysis of migratory behaviour in D & E. Data acquired from 3 independent experiments. n=5, representative video show in supplementary video 10. Correlation with Squared Pearson’s Coefficient. ***p=0.0002; ****p <0.0001. Scale bar = 50 μm. **D,** Quantification of neutrophil migration speed within PNC-bearing epidermis, shows that *myd88*^lo^ immature neutrophils migrate slower than *myd88*^hi^ mature neutrophils. (Mann-Whitney test, N>=48, ****p<0.0001) **E,** Quantification of neutrophil time spend in direct contact with PNCs in time-laps movies, shows that *myd88*^lo^ immature neutrophils remain contact with PNCs for longer than *myd88*^hi^ mature neutrophils. (Mann-Whitney test, N>=48, ****p<0.0001) **F,** The number of differentially expressed genes within each H1-4 neutrophil cluster versus the matching control populations. (Welch’s T-tests. Genes were filtered by expression in at least 10% of cells within at least one of the groups compared, FDR < 0.05 and fold change > |1.25|). **G,** The venn diagram depicts the overlap of upregulated genes within each H1-H4 neutrophil, indicating a shared response regardless of neutrophil maturity. **H,** Heatmap depicts the scaled centred expression of genes upregulated in all H1-H4 neutrophil clusters versus corresponding control clusters. Genes are grouped by biological process.

Recruitment of immature neutrophils to tumours is well documented ^68^, however, their response compared to mature neutrophils remains unclear. To investigate this, we revisited the scRNA-seq data to compare the responses of immature and mature neutrophils following PNC induction. Both immature and mature neutrophils upregulated a similar number of genes versus their maturation matched control clusters (Fig. 7F) (Supplementary Table 5), except the intermediate H2 cluster, which upregulated a great number of cell cycle related genes (Extended Data Fig. 4C). Moreover, immature and mature neutrophils shared many DE genes (Fig. 7G), including those related to the cell cycle, ER protein processing, proteasomal degradation, inflammatory signalling, migration, intracellular signalling, splicing and oxidative phosphorylation (Fig. 7H). Despite the commonality of these responses to PNCs, immature and mature neutrophils maintained distinct developmental states (Extended Data Fig. 4D).

### The initial response of neutrophils to Ras-driven PNCs is tumour-promoting

Neutrophils within tumours can play pro- or anti- tumour roles^69^, therefore, we sought to determine whether their initial response to Ras-driven PNCs was predominately pro- or anti-tumour. Genes typically expressed in immunosuppressive tumour promoting neutrophils, *arg2*^69^ and *ptges3b*^70^, were upregulated in the majority of HRAS neutrophils clusters by 8hpi (Fig. 8A). Classical pro-inflammatory marker *il1b* was downregulated, and *cybb*/*nox2* showed no change (Fig. 8A). This suggests an early skewing towards a tumour promoting immunosuppressive function, which aligns with the observation that PNCs, particularly the PNC-3 cluster, expressed cytokines that are known to drive tumour associated neutrophil development (Fig. 6A, *il1b*, *csf3b* and *tgfb1a* ^69,71,72^).

**Figure 8.**
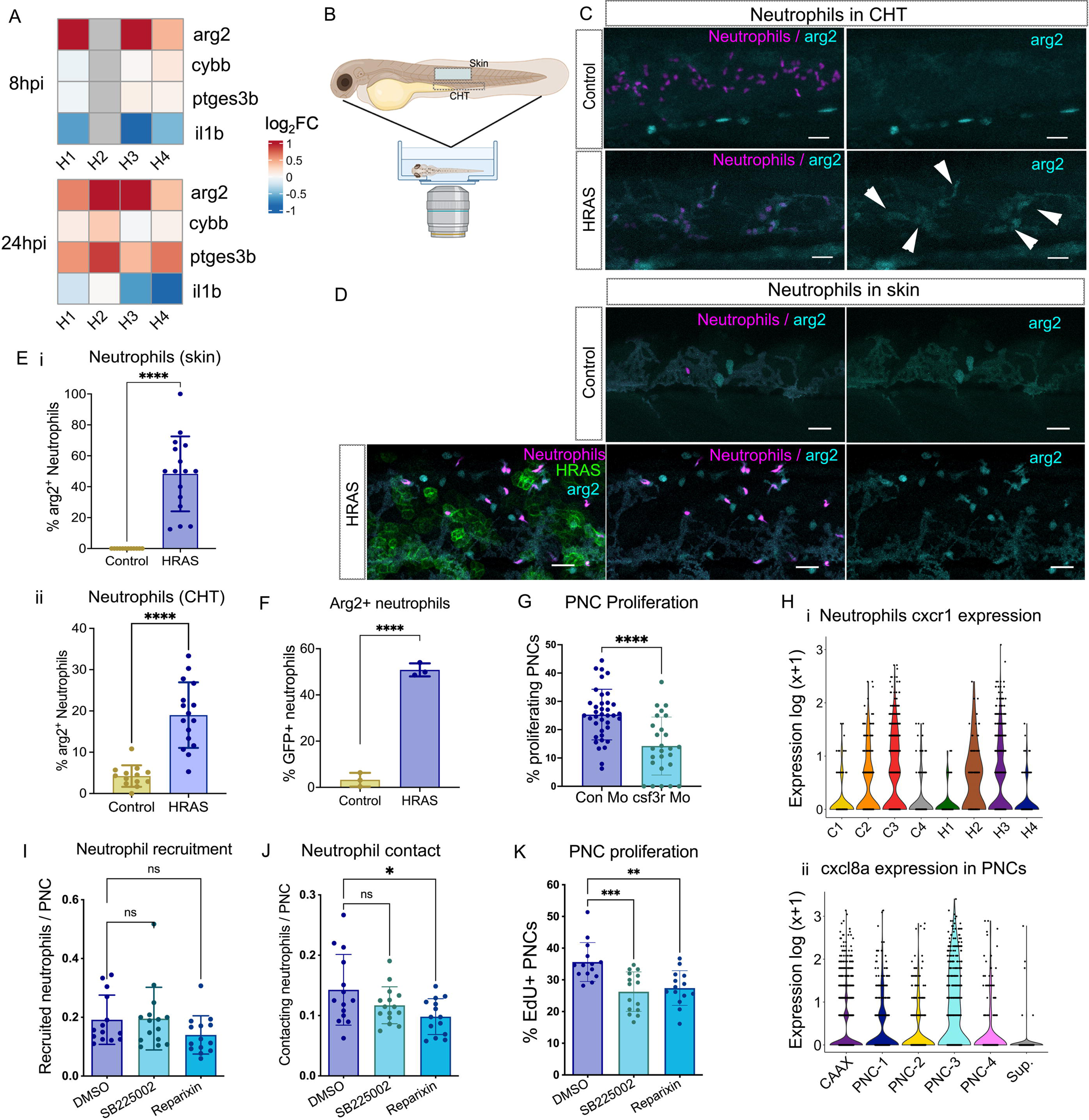
Neutrophils from PNC-bearing larvae express immunosuppressive marker arginase 2 and promote PNC proliferation. **A,** Heatmap depicting log2 fold change of pro-tumour TAN marker genes, *arg2* and *ptges3b*, and pro-inflammatory genes, *il1b* and *cybb* in the neutrophil clusters from PNC-bearing larvae versus their equivalent clusters from control larvae. **B-D**, Confocal live imaging of transgenic *arg2* reporter (turquoise) expressed within neutrophils (magenta) of PNC-bearing larvae versus sibling controls at 24hpi. Scale bar = 30µm. **B,** Schematic showing the regions of the epidermis and the CHT that were imaged. **C,** Images of neutrophils within the CHT of (i) a control larva, and (ii) a PNC-bearing larva. Arrowheads indicate *arg2* reporter expression within neutrophils of the PNC-bearing larva, whilst neutrophils from control larva lacked *arg2* reporter expression. **D,** Images of neutrophils within the epidermis of (i) a control larva, and (ii) a PNC-bearing larva. **E,** Quantification of *arg2* positive neutrophils in: (i) the CHT (p<0.0001), and (ii) epidermis (p<0.0001) (Welch’s t test n>=12). **F,** Quantification of *arg2* reporter expression by flow cytometry analysis of neutrophils from PNC-bearing larvae (HRAS) versus sibling controls at 24hpi (Unpaired t test, n=3, p<0.0001). **G,** The percentage of EdU positive PNCs in larvae injected with *csf3r* morpholino versus control morpholino (Mann-Whitney test, n>=26, p<0.0001). **H**, **i**, Violin plot depicting *cxcr1* expression within neutrophils from the 24hpi scRNA-seq data; **ii,** Violin plot depicting the zebrafish cxcr1 ligand *cxcl8a* expression within PNCs from the 24hpi scRNAseq data. **I,** Quantification of neutrophil recruitment to the epidermis indicated by number of *lyz*:DsRed positive cells normalised by the number of PNCs per field of view. (One-way ANOVA, n>=14, p=0.1796.) **J,** Quantification of direct neutrophil-PNC contact observed by confocal imaging at 24hpi. (One way ANOVA test with Dunnett’s multiple comparisons test, n>=14, p=0.0254.) **K**, Quantification of EdU positive PNCs. (One-way ANOVA test with Dunnett’s multiple comparisons test, n>=14, p=0.0002).

*In vivo* live imaging using an *arg2:GFP* reporter fish^73^, confirmed that neutrophils recruited to the epidermis of PNC-bearing larvae express *arg2:GFP* (Fig. 8D&E). We also observed a significantly increased proportion of *arg2*^+^ neutrophils within the CHT of PNC-bearing larvae (Fig. 8C&E), which is also supported by flow cytometry (Fig. 8F). This provides *in vivo* validation that neutrophils responded to the presence of PNCs systemically and demonstrates that the programming of neutrophils to an “tumour promoting immunosuppressive” began even prior to their recruitment to the local PNC microenvironment.

To establish whether these neutrophils indeed promote PNC development, we used a well-established *gcsf3r* morpholino to generate neutropenic larvae^74^. This led to a substantial decrease in EdU^+^ PNCs at 24hpi (Fig. 8G), indicating that neutrophils promote PNC proliferation. Key neutrophil chemoattractant molecules C*xcl8a* and *cxcl8b* were upregulated within PNCs (Fig. 5A) and are known to function through *cxcr1* and *cxcr2* respectively. *cxcr1* was detected within neutrophils by scRNA-seq (Fig. 8 Hi) whilst its legand cxcl8a is up-regulated in the PNC-3 cluster (Fig8. Hii). Therefore, we used cxcr1/2 inhibitors to block neutrophil recruitment to PNCs. To our surprise, this did not reduce the number of neutrophils recruited to the epidermis (Fig. 8I). However, we did observe a reduction in direct contact between neutrophils and PNCs (Fig. 8J) importantly we saw reduced percentage of EdU^+^ PNCs (Fig. 8K). Thus, we confirmed that neutrophils play a tumour promoting role within the first 24 hours following oncogene activation and identified a role for the cxcl8-cxcr1/2 axis in this process.

## Discussion

Cells carrying oncogenic mutations are found within normal healthy tissues throughout the human body, indicating non-genetic promoting factors are required for tumourigenesis^2,3^. Understanding why some mutant cells form tumours, while others do not, could reveal novel opportunities for cancer prevention. However, few studies have performed detailed analysis during the period between initial oncogenic mutation and tumour formation. Our study is the first to investigate the immediate cell-intrinsic and host responses to oncogenic Ras *in vivo* at the single cell level.

We have found oncogenic Ras is sufficient to unlock cellular plasticity within permissive basal epidermal cells, resulting in rapid dedifferentiation (PNC-1/2) and partial EMT (PNC-3) within 24 hours of induction. Dedifferentiation, evidenced by loss of cellular identity and re-expression of embryonic programmes, likely serves as a gateway to novel cell states required for malignant transformation^20,37,38,75^. Indeed, both scRNA-seq and *in vivo* imaging indicated that dedifferentiation preceded pEMT, with pseudotime predicting a trajectory between these states. We propose that the PNC-3 / pEMT subpopulation is most likely to give rise to tumours, owing to their expression of cancer stem cell markers and drawing a parallel with the high tumour-propagating capacity of hybrid E/M cells^51^.

Whilst lineage tracing would be required to definitively establish that the PNC-3 cells are tumour-initiating cells, we observed that they resemble a lineage-traced cancer stem cell population identified within a murine model of HRAS^G12V^ skin carcinoma^53^. PNC-3 cells and CSCs shared many marker genes, including *lamc2* and *krt18*, the co-expression of which we confirmed *in vivo*. Furthermore, in cross-species integration with patient scRNA-seq data PNC-3 cells mapped to a highly malignant subpopulation located at the invasive edge of tumours^54^, consistent with their high expression of ECM-degrading enzymes and a prior study showing that PNCs within our model degrade their underlying collagen matrix^76^. The PNC-3 signature was also associated with reduced patient survival across various cancer types, particularly those driven by Ras mutations. This further supports the notion that the PNC-3 cellular state contributes to malignancy.

Our findings challenge the idea that malignant properties, such as plasticity, EMT and migration, require multiple mutagenic “hits”. The short time frame in which these state changes occurred strongly implies that the cellular response to oncogenic Ras signalling was determined by pre-existing non-genetic factors intrinsic to the cell of origin. This is especially apparent when comparing basal versus superficial keratinocytes, the latter of which retained their cellular identity and did not undergo cell state changes in response to oncogenic Ras. Superficial keratinocytes are terminally differentiated, whilst the basal population consists of stem and progenitor cells, suggesting that stem cells are more susceptible to oncogene-induced plasticity. This aligns with previous claims that CSCs originate from tissue stem cells^77^ and may explain early murine studies of skin tumourigenesis in which oncogenic Ras drove tumour formation within the basal layer of the epidermis, but not the suprabasal layers^7,78^. Interestingly, even within our basal-derived PNC population, some cells progressed to pEMT (PNC-3) whilst others differentiated to suprabasal fate (PNC-4). It is possible that these cells originated from stem and progenitor cells respectively, since basal progenitors are committed to differentiation^79,80^.

Factors within the local microenvironment can also influence tumourigenesis, such as inflammation^9^. A recent study demonstrated that extrinsic inflammation unlocks a malignant cell state within Ras mutant cells^81^. Our live imaging studies showed a rapid myeloid response, wherein recruited neutrophils appeared to preferentially interact with mesenchymal-like PNC-3 cells. Neutrophils were programmed to an *arg2* ^+ve^ state and promoted the proliferation of PNCs. This could be partially attenuated by reducing PNC-neutrophil interactions via blocking *il8-cxcr1/2* axis. Furthermore, a previous zebrafish study using *cxcr1/2* inhibitors demonstrated that intrinsic neutrophilic inflammation promotes EMT within Ras mutant skin PNCs^82^. Strikingly, our transcriptomic data revealed PNC-3 cells were a major source of chemokines and cytokines known to instigate pro-tumour myeloid responses. Together, these data suggest that these emergent pre-malignant cells reinforce their own cellular state by creating a positive feedback loop with tumour-promoting neutrophils.

Amongst the cytokines enriched within the PNC-3 cluster was *csf3b*, the ortholog of G-CSF, a systemic signal that triggers premature release of neutrophils from haematopoietic tissues, facilitating the recruitment of immature pro-tumour neutrophils to tumours^65,66,68^. Akin to a recent study of tumour associated neutrophils (TANs) within the tumour microenvironment^83^, both immature and mature neutrophils were recruited to PNC bearing epidermis and underwent similar transcriptomic responses. However, instead of convergent to a common state as in the TANs, PNC recruited neutrophils maintained their developmental identity and displayed different migratory behaviours. Functional differences between immature and mature neutrophils have also been reported in *ex vivo* assays^59^. Our zebrafish model would be useful for future research on this subject.

In summary, our work demonstrates that oncogenic Ras can unlock cellular plasticity from the inception of preneoplastic development, creating cellular states absent in normal tissue but resembling malignant cells found within established tumours. Malignant features were only acquired by permissive cell subpopulations, via dedifferentiation and partial EMT. These cells resembled cancer stem cells and were the primary instigators of intrinsic inflammation, creating a tumour-promoting microenvironment via neutrophil reprogramming. Our focus upon the first 24 hours of Ras-induced preneoplastic development reveals the rapid onset of both cellular plasticity and inflammation, raising far-reaching implications for early detection and cancer prevention.

## Materials and methods

### Transgenic Zebrafish strains

The following zebrafish lines were generated in the Feng laboratory using Tol2 mediated transgenesis^24^: Tg(UAS::EGFP-HRAS^G12V^; cmlc2::EGFP)^ed203^, Tg(UAS::mCherry-HRAS^G12V^; cry::CFP)^ed205^, Tg(UAS::EGFP-CAAX; cmlc2::EGFP)^ed204^, Tg(UAS::BFP-HRAS^G12V^; cry::CFP)^ed209^ & Tg(UAS::mCherry-CAAX; cry::CFP)^ed206^. As were Tg(krtt1c19e::KalTA4-ER^T2^; cmlc2::EGFP)^ed201 76^,Tg(lyz: lifAct-mTuquoise2a)^ed211^ and Tg(lyz::NLS-mScarlet)^ed229 84^. The arginase reporter line, TgBAC(arg2:EGFP)^sh571^ and Tg(myD88:DsRed) was reported previously^73,67^.

### Zebrafish maintenance and breeding

Adult zebrafish were maintained in the Bioresearch & Veterinary Services (BVS) Aquatic facility in the Queen’s Medical Research Institute, the University of Edinburgh. Housing conditions were described in the Zebrafish Book^85^. All experiments were conducted with local ethical approval from The University of Edinburgh and in accordance with UK Home Office regulations (Guidance on the Operations of Animals, Scientific Procedures Act, 1986) under the authority of the Project Licence PEE579666.

### 4-Hydroxy-tamoxifen induction and drug treatment

Induction of HRAS^G12V^ expression was performed as previously described^27^. In brief, 0.3× Danieau’s solution containing 5 µM 4-Hydroxy-tamoxifen (4-OH-tamoxifen, SIGMA) and 0.5% DMSO was used to submerge larvae from 52 or 72hpf onward. Drug treatments were carried out between 8hpi – 24hpi, when fluorescent HRAS^G12V^ first became visible, using SB225002 1µM or Reparixin 1µM.

### Morpholino knockdown of csf3r

A 1 nl bolus of 1 mM *csf3r* morpholino (GeneTools, 5’-GAAGCACAAGCGAGACGGATGCCAT-3’)^74^ was injected into the yolk of each egg. 1 mM human β-globin morpholino (GeneTools, 5’-CCTCTTACCTCAGTTACAATTTATA-3’) was used as control.

### EdU (5-ethynyl-2ʹ-deoxyuridine) incorporation and staining

For detailed protocol refer to protocol.io^86^. In brief, 2nl of EdU solution (10mM) was injected into the yolk of individual larva. Larvae were fixed at 2.5 hours post injection, with 4% PFA for 30 minutes. EdU staining was carried out for 30 minutes using Click-iT Plus Edu Alexa FluorTM 647 Imaging Kit (Thermo Fisher Scientific, C10640).

### Wholemount immunostaining for Cleaved-Caspase 3

Neutrophil apoptosis was assessed using wholemount anti cleaved-Caspase 3 immunostaining. In brief, Larvae were fixed in 4% PFA for 2 hours at room temperature followed MeOH permeabilization at -20°C. Larvae were re-hydrated at room temperature then blocked for 2 hours before overnight incubation with the primary antibody (Purified Rabbit Anti-active caspase 3 (BD 559565) 1:200 in Block solution) at +4°C. On day 2, larvae were washed 6 x 20 min with 0.1%PBT at room temperature, followed by 2 hours incubation with secondary antibody (Alexa Fluor 633 goat anti-rabbit IgG (H+L) (Invitrogen A21071) 1:250 in block solution). The larvae were washed 3 x 15 min in 0.1% PBT at room temperature, then mounted in Citi Fluor for confocal imaging.

### HCR RNA fluorescence in situ hybridisation (HCR RNA-FISH)

The HCR *in-situ* probes for *lamc2* (XM_003197884.5), *cldna* (NM_131762.2) and *krt18a.1* (NM_178437.2) were synthesised by Molecular Instruments. The HCR protocol was performed according to the manufacturer’s instructions with the exception of proteinase K treatment, which was reduced to 8 μg/ml for 15 mins in order to preserve the epidermis.

### Total RNA extraction from whole larvae

Multiple larvae with the desired phenotype (40-50 larvae per group) were transferred to 1 ml TRIzol reagent (Invitrogen) and stored overnight at -80°C. Upon thawing samples were homogenised by repeatedly passing through a 23-gauge needle (BD Microlance). Total RNA was isolated with chloroform extraction followed by isopropanol precipitation. After final 70% Ethanol wash, pellet was air-dried and resuspended in Nuclease-Free Water (Invitrogen, AM9930).

### cDNA synthesis from total RNA

cDNA synthesis was achieved using the iScript™ Advanced cDNA synthesis Kit for RT-qPCR (Bio Rad, 1725037). As per manufacturer’s instruction, 4 μL 5x iScript Advanced Reaction Mix and 1 μL iScript Advanced Reverse Transcriptase were added to 15 μL isolated RNA. Resulting reaction mixture was incubated at 46°C for 20 minutes and then at 95°C for 1 minute, in a Mastercycler Nexus Gradient GSX1 Thermal Cycler (Eppendorf).

### Generation of single cell suspension from zebrafish larvae

Detailed protocol can be found in protocol.io^87^. In brief, lots of 50 larvae were suspended in 2ml Media A: HBSS -Mg, -Ca, + phenol red (Gibco, 11530476), 1M HEPES (Sigma Aldrich, H3375), 25mM D-Glucose (Sigma Aldrich, G8644), 2% Sterile Goat Serum (Life Technologies, 16210064). Media A was supplemented with 2.5 mg/ml Collagenase IV (Life Technologies, 17104019). Larvae were transferred to a 12 well plate and incubated at 28.5 °C, with shaking at 100 rpm for 5 minutes, followed by mechanical disruption, i.e., rapid pipetting with a P1000 Gilson pipette for 30 seconds. After 3 rounds of incubation and rapid pipetting, digests from 3 wells were pooled into a Falcon tube. The digests were then centrifuged at 4°C for 5 minutes at 500g and the pellet was resuspended in 500μl of Media A.

### Flow Assisted Cell Sorting of keratinocytes, PNCs and myeloid cells

Larval digests were passed through a 40μM mesh and DAPI was added for live/dead detection (1:1000). Keratinocytes and PNCs were selected by the fluorescence of the mCherry-CAAX and mCherry-HRAS^G12V^ transgenes respectively. Myeloid cells were selected by fluorescence from the mpeg1.1:EGFP reporter. Autofluorescent cells were excluded by gating out cells with fluorescence in both B 525/50 and B 695/40 filters. Both mCherry and EGFP positive cells were sorted into a single 1.5ml Eppendorf containing 50μl of Media A supplemented with 10% Sterile Goat Serum at 4°C.

### Flow cytometry analysis of transgenic *arg2* reporter

Single cell suspensions from dissociated larvae were stained with 1:1000 DAPI (Thermofisher, #62248) to indicate cell viability and flow cytometry was carried out at the QMRI Flow facility using a BD 5 Laser LSR Fortessa cell analyser. The gating strategy used is shown in supplement material (extended data Fig. 5). All data was analysed in FlowJo v10.8.1 software (BD Life Sciences).

### cDNA synthesis from FACS sorted cells following the adapted Smart-seq2 protocol

FACS sorted HRAS or CAAX populations were collected directly into Eppendorf tubes containing 4 µL 0.2% Triton X and RNAase inhibitor (Thermo Fisher Scientific) at 1,000 cells per tube. The smart-seq2 protocol was used to prepare the samples for cDNA synthesis^89^, cDNA quality was assessed using a LabChip GX machine.

### Quantitative PCR

Gene expression levels were measured on a LightCycler 96 system using the LightCycler 480 SYBR Green I Master mix (Roche). The primer sequences were the following: *arpc5a* forward 5’-AGTGCTGAAGAATCCACCCA-3’ and reverse 5’-ATCGCTAGCCTTGAAAGAGC-3’; *snail2* forward 5’-GACGCACAGAGTCGGAAATC-3’ and reverse 5’-TCTAATGTGTCCCTGCAGCA-3’; *mmp9* forward 5’-ACCCTCGATTCACTGATGCA-3’ and reverse 5’-CTCCAGTAGAAGCTCTCCCG-3’; *itgb1b* forward 5’-TGCGACAACTTTAACTGCGA-3’ and reverse 5’-TGCCGGTGTAGTTAGCATCA-3’, *itga5* forward 5’-TGTGTAATGTTGGCCTGTTGG-3’ and reverse 5’-TGTAACTAGCATGGCACTCC-3’; *itga3* forward 5’-TGTGATCAGCCAGCGAGA-3’ and reverse 5’-CATCTGAAGGTTGCTCTCGC-3’. *cnppd1* forward 5’-TGGCGATTTGTTTGACGAGC-3’ and reverse 5’-TCACTCTCTCTGACCACTGC was used as the house-keeping gene.

### 10X single cell RNA-sequencing

For each sample, 60,000 cells were sorted by FACS and washed by centrifugation at 500g for 5 minutes at 4°C, followed by replacement of the supernatant with 20μl HBSS. A Countess 3 automated cell counter was used to calculate cell density, and 7,000 live cells were loaded on to each channel of the Single Cell chip (10X Genomics). Single cells were encapsulated in droplets using the Chromium Controller (10X Genomics). Sequencing libraries were prepared using the Chromium Single Cell 3’ kit v3 as per the manufacturer’s protocol. The LabChip GX Touch Nucleic Acid Analyser was used to quantify the concentration and quality of the libraries. Libraries were sequenced on a single Illumina Novaseq S3 flow cell with 300-400 million raw reads per sample.

### Analysis of zebrafish single cell RNA-sequencing data

Single cell RNA-seq data was demultiplexed and aligned using the CellRanger pipeline (10x Genomics Cell Ranger 3.1.0) and the Ensembl GRCz11.101 reference genome. Cells with fewer than 500 genes detected and and/or greater than 12 percent mitochondrial reads were removed. Genes that were expressed within less than 2 cells were also removed. Ambient RNA was detected and removed using SoupX^90^. To detect doublets, three methods were used: doubletCells from scran, Scrublet^91^ and DoubletDecon^92^. For the former two methods, a cut-off score was determined by inspecting a histogram to identify an outlying population with high doublet scores. Cells that were called doublets by at least two of these methods were discarded.

SCTransform^93^ was used to perform normalisation, scaling, selection of highly variable genes and regression of cell cycle genes. Genes related to “S” and “G2M” phases of the cell cycle were identified using the Gene Ontology data base for Danio rerio. Genes associated with the following GO terms were assigned to the “S phase” gene list: “DNA replication”, “DNA repair” and “mitotic DNA replication”. Genes associated with the following GO terms were assigned to the “G2M phase” gene list: “chromosome segregation”, “mitotic nuclear division” and “mitotic cytokinesis”. Genes which overlapped between the two lists were removed. The following genes were added manually to the “S phase” gene list: *ccnd1, ccnd2a, ccnd2”, ccnd3, ccne1, ccne2, meis1b, ccne1, ccne2, ccna1, ccna2, cdc6, mcm2, mcm3, mcm4, mcm5, mcm6, mcm7, mcm10, orc1,* and *orc6.* The following genes were added manually to the “G2M phase” gene list: *ccna1, ccna2, foxm1, skp2, cdk1, ccnb1, ccnb2, ccnb3, mki67, top2a, oip5, nusap1, kif20a, kif23, racgap1, sgo1* and *anln*. Cell cycle scoring was performed in Seurat.

Where required, samples were integrated using Harmony^94^ prior to Louvain clustering. To discern cell types, a priori markers were inspected and further confirmed by inspection of DE genes within each cluster. Differential abundance of the various skin cell types between conditions was tested using EdgeR. The estimateDisp function was used to estimate negative binomial dispersion, the quasi-likelihood dispersion was estimated by glmQLFit and testing was performed by glmQLFTest. Pseudobulk differential expression analysis were performed on cells of equivalent type between the CAAX and HRAS samples using EdgeR. Raw counts were summed across groups and normalised. The negative binomial dispersion of each group was calculated with estimateDisp, the quasi-likelihood dispersion was estimated using glQLFit and testing was carried out with glmQLFTest.

For detailed analysis of keratinocytes/PNCs, neutrophils and macrophages, the data was subset by cell type and re-normalised prior to sub-clustering. Single cell differential expression analyses between clusters were performed using the findMarkers function from scran. For permissive differential expression analysis, Welch’s t-tests were used to compare each cluster versus the sample. Wilcoxon Rank Sum Tests were performed between each pairwise combination of clusters to narrow down specific genes. Results were filtered for genes that were expressed within >10% cells per cluster of interest, and FDR < 0.05. To identify the optimal markers of the PNC clusters, three tests were used with the following thresholds: pairwise Welch’s t-tests (fold-change > 2), pairwise Wilcoxon Rank Sum tests (are under the curve > 0.8), and pairwise Binomial tests (fold-change > 2), and genes were filtered for expression with > 75% of cells within the cluster of interest.

Gene Set Enrichment Analysis (GSEA) was carried out using software (version 4.2.2) from The Broad Institute to test for enrichment of Biological Processes from the *Danio rerio* Gene Ontology database (FDR < 0.1). Over-representation analysis was performed using a hypergeometric test with R package fgsea. The gene sets for “immature Basal Epidermis” and “Mature Basal Epidermis” were derived from a scRNA-seq time course of zebrafish embryonic development^39^. Nascent epidermal cells (10pf) were compared with developed epidermal cells (24hpf) using Welch’s T-test (FDR < 0.05, fold change > 1.25 and detected in > 10% of cells). The gene sets for “Upregulated by *tp63*” and “Downregulated by *tp63*” were taken directly from Santos-Pereira et al. 2019, who performed transcriptomic comparison between *tp63* knockout versus wildtype zebrafish larvae. The gene sets for “Basal Keratinocyte”, “Superficial Keratinocyte” and “All Keratinocyte” were taken directly from Cokus et al. 2019, who performed a transcriptomic comparison of the basal keratinocytes and superficial keratinocytes of 72dpf zebrafish larvae in comparison to all keratinocytes and non-keratinocytes.

Trajectory analysis was performed using Monocle 3^95^. The raw counts from the PNCs of the HRAS samples were re-normalised and batch corrected, followed by dimensionality reduction and clustering by the standard pre-processing guidelines for Monocle 3, prior to ordering and scoring by pseudotime. The cells within the 8hpi samples that most resembled their basal keratinocyte cells of origin were selected as the root.

### Cross-species integration of zebrafish and human scRNA-seq datasets

Human orthologues of zebrafish genes were acquired from the Alliance of Genome Resources and multi-mapping genes were excluded from downstream analysis. ScRNA-seq data from human cutaneous squamous cell carcinoma tumours and healthy skin was obtained from Ji et al. 2020^54^ (GSE144240) and merged with the scRNA-seq data from keratinocytes and PNCs published herein. LIGER^96^ was utilised to directly integrate the two data sets as follows. Firstly, the merged datasets were re-normalised, scaled and highly variable genes selected according to LIGER default settings. Joint matrix factorisation was performed (k = 15) and the resulting factors were manually inspected. Factors dominated by cell cycle genes and ribosomal genes were excluded from downstream analysis. Finally, the datasets were integrated by quantile normalisation and jointly clustered using Louvain community detection.

### Confocal imaging & analysis

Confocal fluorescent images were acquired with Leica confocal SP8, using HCX PL APO 40x or 25X water immersion objective, or HC PL APO 20x CS2 dry objective (overnight time-laps). Whole body neutrophil imaging were done with Opera Phenix Plus (Revvity) microscope using 20x water objective (NA 1.0). Appropriate lasers and collection gates were selected according to excitation and emission wavelength for each fluorophore. Data analyses were carried out using IMARIS 9.0. Percentage of TUNEL positive cells within EGFP positive cells were manually counted by inspecting each optical section of image Z-stacks. Neutrophil numbers and positioning in the whole larvae and percentage of EdU^+ve^ neutrophils were quantified in 3D using Harmony 5.2 software (Revvity). Neutrophil and macrophage footprint analysis were carried out using Volocity 6.3 (Perkin Elmer) tracking fluorescent object function. Time-lapse videos were exported from Volocity, Imaris 9.0 or ImageJ.

### PanCancer RNA-seq analysis

Finalised RNA-seq and mutation data from the Pan-Cancer Atlas, generated by the Cancer Genome Atlas (TCGA), were obtained from the Genomic Data Commons portal (https://gdc.cancer.gov/)(The Cancer Genome Atlas Program (TCGA) - NCI). RAS mutation status was then inferred by non-silent mutation of KRAS, NRAS, or HRAS at residues G12, G13, and Q61, which are believed to account for 98% of all RAS mutations^98^. In order to assess expression of the PNC-3 cluster genes within the context of mutation status, the collective expression of the top 30 differentially expressed genes within the signature was quantified as a single metric via gene set variation analysis (GSVA)^99^. Survival statistics were calculated and plotted for cancer-types with >10% RAS mutation frequency using the survival and survminer R packages, with patients stratified by high/low expression using thresholds calculated via the SurvivALL package^100^.

### Statistical analysis

Appropriate parametric or non-parametric tests to determine differences in mean or rank were carried out using Graphpad Prism 9.1.4 software (San Diego, USA). Degrees of freedom were calculated from the number of individual larvae measured. Graphs depict mean ± standard deviation (SD) unless otherwise stated. Null hypotheses were rejected when *p* < 0.05.

## Acknowledgement

The authors thank the technical support from the Confocal Advanced Light Microscopy facility in the Queen’s Medical Research Institute & in the Institute For regeneration and Repair. The High Content Screening Facility in the Institute for Regeneration and Repair and the BVS Aquatic Facility Units at the University of Edinburgh. The authors thank Dr. Bartlomiej Waclaw for assistant in performing imaging data analysis. The results shown in Figure 6 G are based upon data generated by the TCGA Research Network: https://www.cancer.gov/tcga. The authors acknowledge fundings from Wellcome Trust Sir Henry Dale Fellowship to Y.F. (100104/Z/12/Z); Cancer Research UK Early Detection Award to Y.F. (C38363/A26931); A.M.E. was supported by a Medical Research Council PhD Studentship (MR/N013166/1); H. M. was supported by a postdoctoral fellowship from the Sigrid Jusélius Foundation. P.M.E. is funded by a Sir Henry Dale Fellowship, jointly funded by the Wellcome Trust and the Royal Society (Grant Number 105570/Z/14/A); C. G. H & R. C are funded by Worldwide Cancer Research (19-0238), Wellcome Trust ISSF 3 (UoE) and LifeArc-CSO; N.C.H is funded by a Wellcome Trust Senior Research Fellowship (219542/Z/19/Z).

---

**Extended Data Figure 1.**
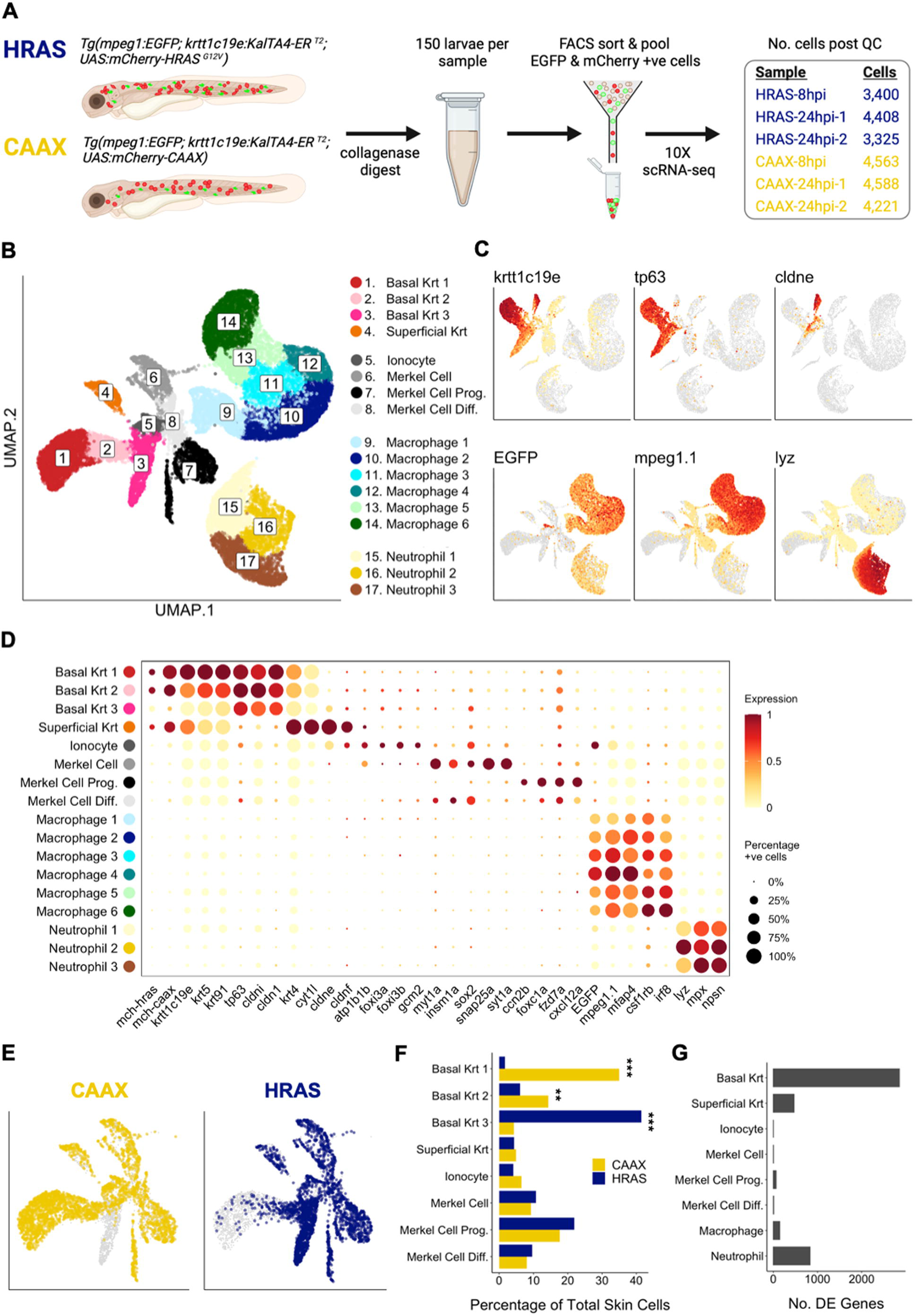
Single cell RNA sequencing of myeloid and epidermal cells. **A,** Schematic showing the workflow for scRNA-seq of myeloid and epidermal cells from Tg(mpeg1:EGFP; krtt1c19e:KALTA4-ER^T2^; UAS:mCherry-HRAS^G21V^) larvae and Tg(mpeg1:EGFP; krtt1c19e:KALTA4-ER^T2^; UAS:mCherry-CAAX) larvae, referred to as CAAX and HRAS larvae respectively. Larvae were induced at 3 dpf with 4-OHT and subjected to digestion with collagenase to form a single cell suspension at either 8 or 24hpi. 150 larvae were pooled per sample. Epidermal cells (mcherry) and myeloid cells (EGFP) were sorted by their fluorescence and pooled prior to library preparation with the 10X Chromium platform. Following initial data processing and elimination of low-quality cells and doublets, 3,000-4,500 high quality cells remained per sample. **B,** UMAP depicts the clustering of all high-quality cells from each sample. **C,** UMAPs depict the normalised expression of epidermal and myeloid markers: *krtt1c19e* (all keratinocytes), *tp63* (basal keratinocytes), *cldne* (superficial keratinocytes), EGFP (all myeloid cells), *mpeg1.1* (macrophages), *lyz* (neutrophils). **D,** Dot plot showing the scaled expressed of an extended panel of transgenes and cell type markers. All keratinocytes = *krtt1c19e*. Basal keratinocytes = *krt5*, *krt9*, *tp63*, *cldni*. Superficial keratinocytes = *krt4*, *cytl1*, *cldne*, *cldnf*. Ionocytes = *atp1b1b*, *foxi3a*, *foxi3b*, *gcm2*. Merkel cells = *myt1a*, *insm1a*, *sox2*, *snap25a*, *syt1a*. Merkel cell progenitors: *ccn2b*, *foxc1a*, *fzd7a*, *cxcl12a*. Macrophage = *mpeg1.1*, *mfap4*, *csf1rb*, *irf8*. Neutrophils = *lyz*, *mpx*, *npsn*. **E,** UMAPs depicting epidermal cells only, from CAAX and HRAS samples. **F**, The percentage of total skin cells assigned to each skin cell cluster at 24hpi, tested for differential abundance in the CAAX versus HRAS samples. Shows that populations of basal keratinocytes are altered, whilst superficial keratinocytes, ionocytes and merkel cells remain stable between conditions (*** indicates P < 0.001, ** indicates P < 0.01). **G,** Differential expression results for each cell type within the HRAS samples, compared to the corresponding CAAX samples at 24 hpi. Shows that basal keratinocytes have a dramatic transcriptomic response, neutrophils and superficial keratinocytes have a modest response, and macrophages, ionocytes and merkel cells have minimal to no response (FDR < 0.05, fold-change > |1.5|).

**Extended Data Figure 2.**
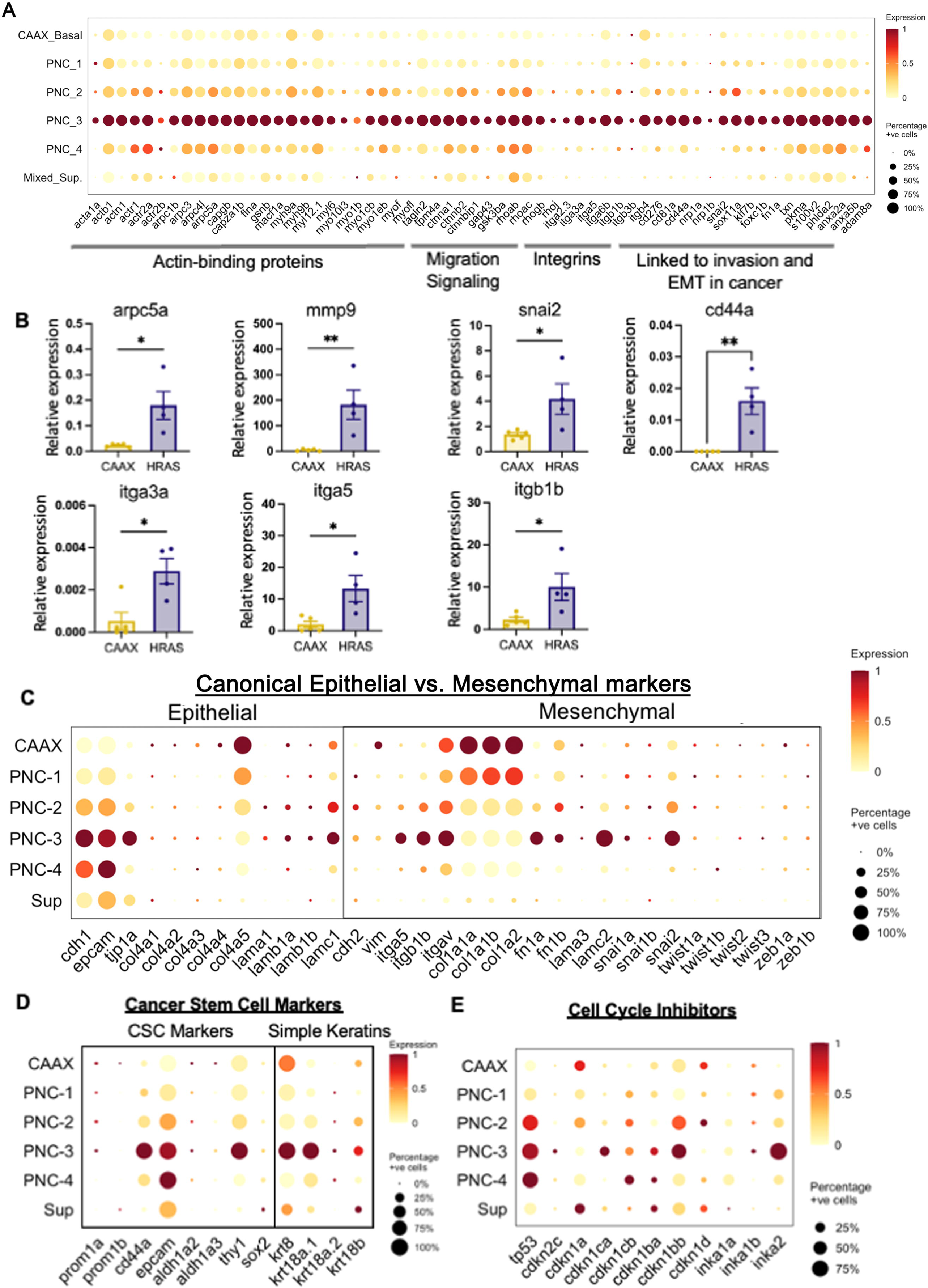
The PNC-3 cluster expresses a combination of epithelial and mesenchymal genes, in addition to selected cancer stem cell markers. **A,** Dot plots depicting the scaled and centred expression of PNC-3 enriched genes related to EMT. **B,** qPCR shows the upregulation of EMT related genes and cancer stem cells marker cd44a in FACS sorted PNCs versus control keratinocytes (Unpaired t test, ** indicates P < 0.01, * indicated P <0.05). **C-D,** Dot plots depicting the scaled and centred expression of: **C,** canonical epithelial and mesenchymal markers; **D,** Cancer stem cell markers; **E,** cell cycle inhibitors.

**Extended Data Figure 3.**
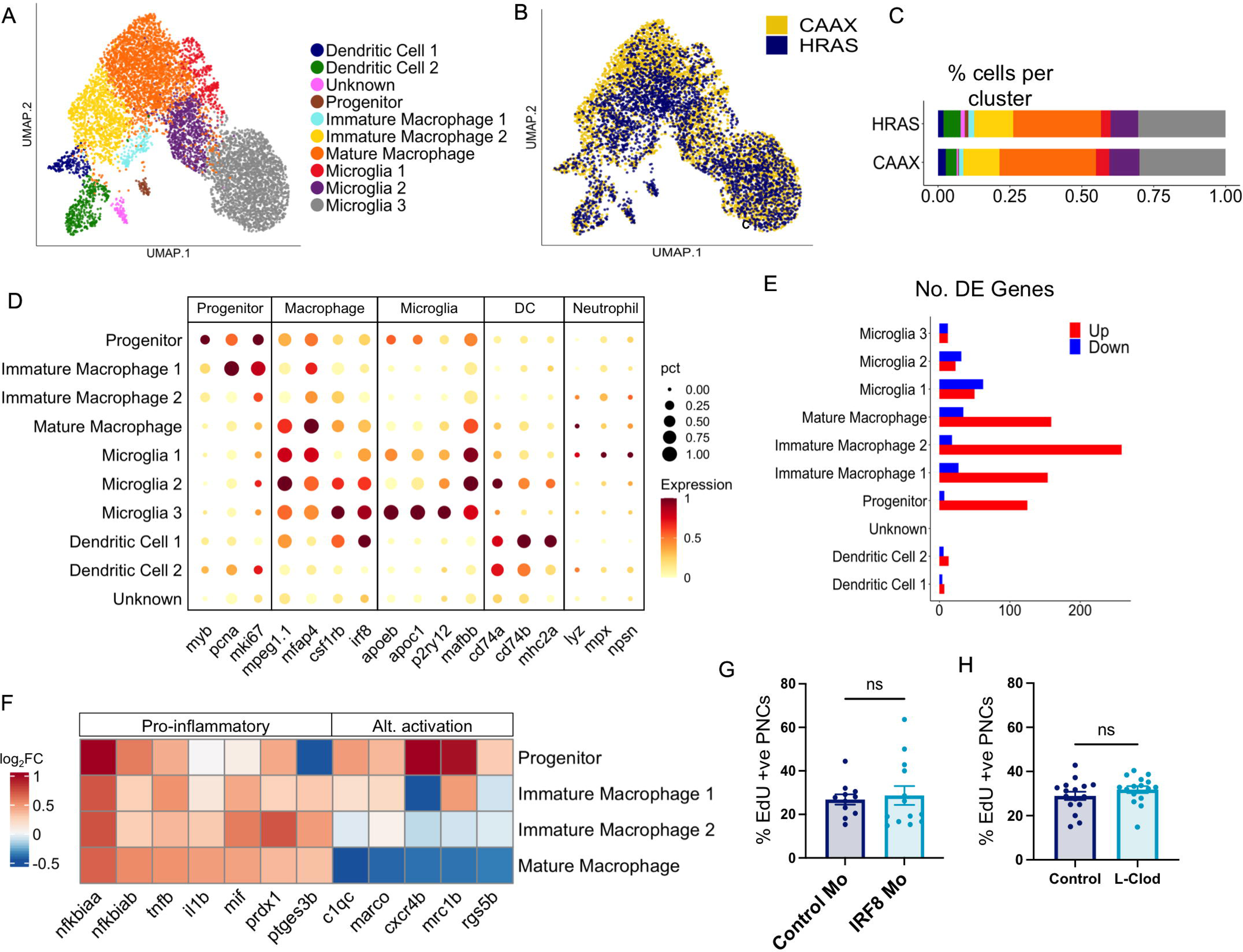
The macrophage response to PNC initiation is minimal and has no effect on PNC proliferation at 24hpi. **A,** UMAP depicts clusters of macrophages, dendritic cells and microglia from 24hpi scRNA-seq samples. **B,** UMAP depicting cells from CAAX and HRAS larvae, shows a high degree of overlap between conditions. **C**, Quantification of the percentage of cells per cluster from CAAX and HRAS larvae shows minimal change in cluster sizes between conditions. **D,** Dot plot showing the expression of macrophage, microglia, dendritic cell and neutrophil markers. **E,** The number of differentially expressed genes identified between CAAX and HRAS samples for each cluster depicted. (Welch’s T-tests. Genes were filtered by expression in at least 10% of cells within at least one of the groups compared, FDR < 0.05 and fold change > |1.25|). **F,** The log2 fold change of genes related to the pro-inflammatory or alternative activation of macrophages. **G,** Quantification of PNC number (i) and the percentage of EdU positive PNCs (ii) from PNC-bearing larvae lacking macrophages due to *irf8* Morpholino injection versus control Morpholino. n=14 (irf8 Mo), n=11 (control Mo), Welch’s t-tests, ns = P >0.05. **H,** Quantification of PNC number (i) and the percentage of EdU positive PNCs (ii) from PNC-bearing larvae depleted of macrophages by injection with Liposome Clodronate (L-Clod). n=16, Welch’s t-tests, ns = P >0.05. **I,** Fluorescence images of macrophages (mpeg1.1:EGFP) in CAAX and HRAS larvae. Boxed area indicates the central haematopoietic tissue (CHT). **J,** Quantification of the mean fluorescent area as a proxy for macrophage abundance within: (i) the whole image, (ii) the CHT, and (iii) the periphery, i.e., outwith the CHT.

**Extended Data Figure 4.**
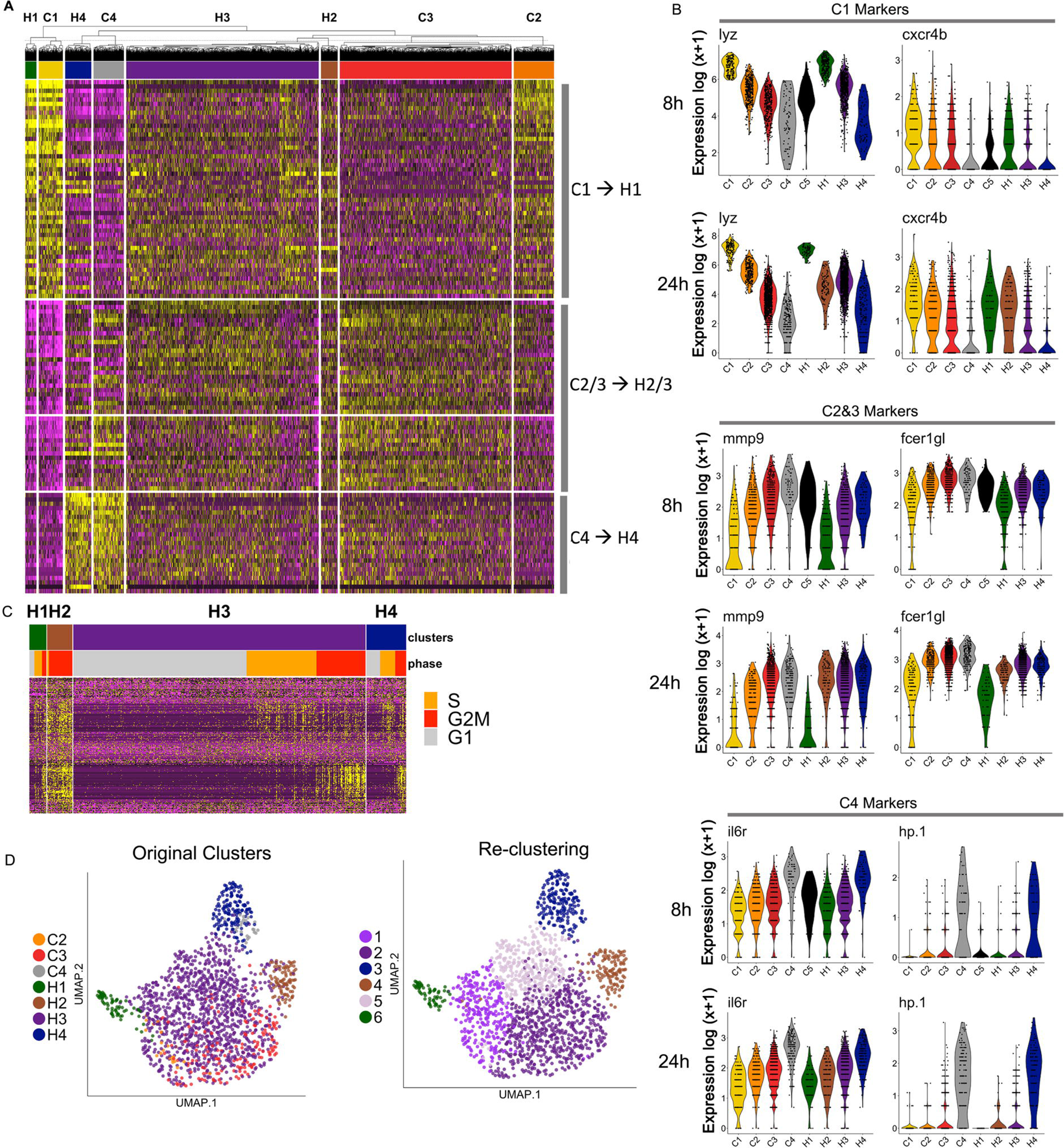
Identification of matched control and activated neutrophil clusters from CAAX and HRAS larvae. **A,** Heatmap depicts the top 50 enriched genes within each control neutrophil clusters (C1-4) versus all remaining CAAX neutrophils. Hierarchical clustering of cell groups shows each neutrophil cluster from the PNC-bearing larvae (H1-4) paired with its matched control cluster, i.e., C1 and H1, C2/3 and H2/3, C4 and H4. (Non-unique genes are included in the cluster in which they first appear in numerical order. Gene expression values are scaled and centred around zero. Differential expression was determined by Welch’s T-test between the cell of a given cluster and the cells from all remaining neutrophil clusters. Genes were filtered by detection in at least 10% of the cells within the given cluster and FDR < 0.05, then ranked by fold-change). **B,** Violin plots showing the normalised expression of genes enriched within C1, C2/3 and C4 clusters. Shows the correspondence between matched CAAX and HRAS neutrophil clusters at both 8 and 24hpi. **C,** Heatmap depicting the 324 genes significantly upregulated within the H2 cluster only. Cells are ordered by cell cycle phase, demonstrating that the majority of these genes correlated with cell cycle phase and were also expressed within proliferative cells of other neutrophil clusters. **D,** Neutrophils from the HRAS samples only were subset and subjected to re-clustering. UMAPs show (i) the original clusters, (ii) the result of re-clustering. This demonstrates that neutrophils from HRAS samples robustly cluster by their developmental state.

**Extended Data Figure 5.**
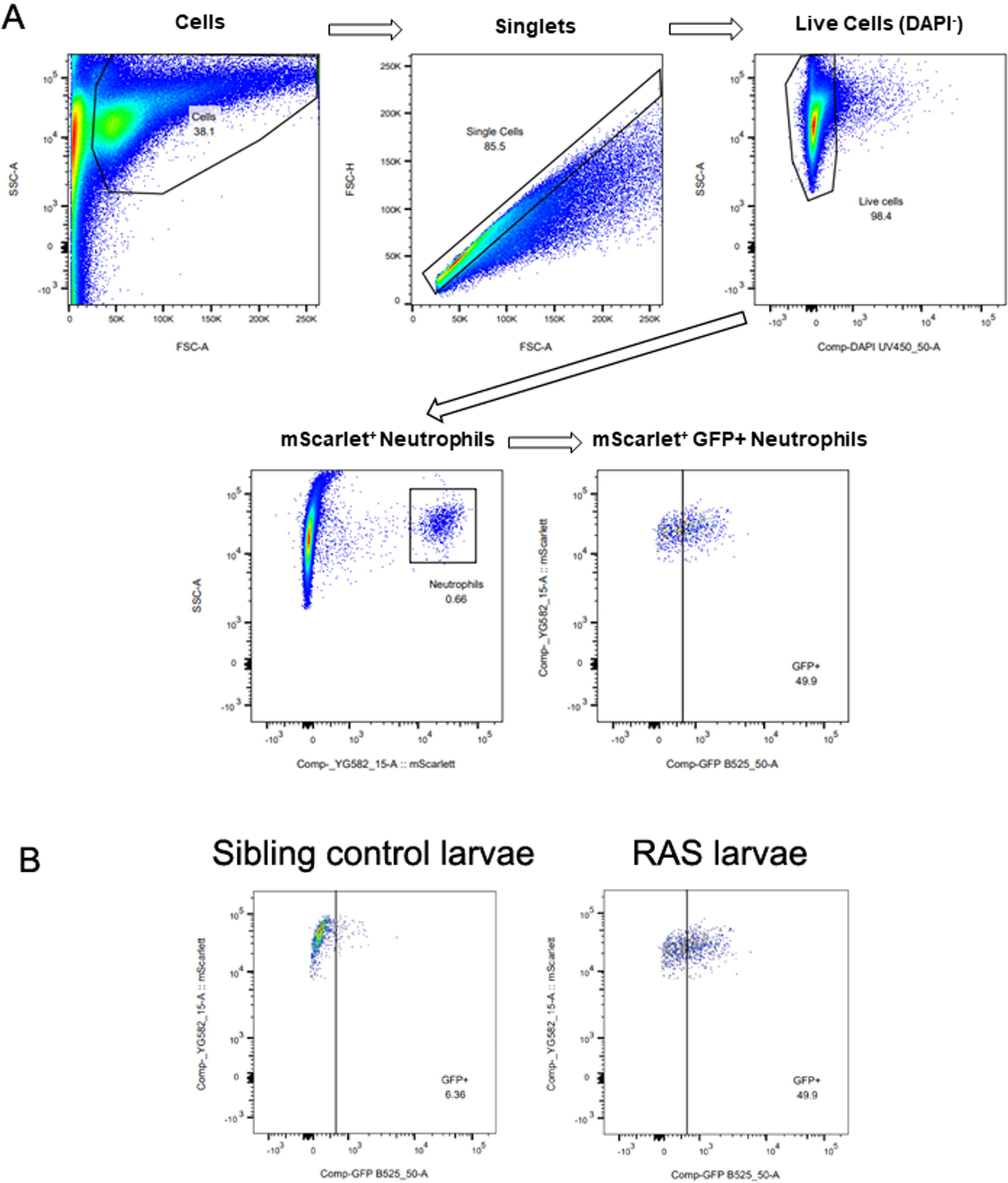
Gating strategy for FACs analysis of *arg2*:eGFP reporter activity in neutrophils. **A**, the gating strategy. **B,** example plots for *arg2*:GFP positive cells within neutrophils in sibling control larvae vs RAS^+ve^ PNC bearing larvae.

**Supplementary Table 1)** Differential expression results for keratinocyte and PNC clusters. Differential expression was carried out in a permissive manner to capture both similarities and differences between clusters, i.e., Welch’s T tests between each cluster versus all other keratinocytes and PNCs.

**Supplementary Table 2)** Pseudobulk differential expression results for each 24hpi PNC cluster versus CAAX control group.

**Supplementary Table 3)** PNC-3 marker gene list for survival analysis using the TCGA database.

**Supplementary Table 4)** Genes differentially expressed within each neutrophil cluster at 24hpi. Welch’s T tests were performed between each pairwise combination of clusters and summary statistics were calculated to identify genes specific to each cluster.

**Supplementary Table 5)** Differential expression results for each HRAS neutrophil cluster versus maturation matched control groups at 24hpi were calculated by Welch’s T tests.

## References

1. Li, S., Balmain, A. & Counter, C. M. A model for RAS mutation patterns in cancers: finding the sweet spot. Nat Rev Cancer 18, 767–777 (2018).

2. Martincorena, I. et al. High burden and pervasive positive selection of somatic mutations in normal human skin. Science (1979) 348, 880–886 (2015).

3. Kakiuchi, N. & Ogawa, S. Clonal expansion in non-cancer tissues. Nat Rev Cancer 21, 239–256 (2021).

4. Goerttler, K. & Loehrke, H. Diaplacental carcinogenesis: initiation with the carcinogens dimethylbenzanthracene (DMBA) and urethane during fetal life and postnatal promotion with the phorbol ester TPA in a modified 2-stage Berenblum/Mottram experiment. Virchows Arch A Pathol Anat Histol 372, 29–38 (1976).

5. Balmain, A. Peto’s paradox revisited: black box vs mechanistic approaches to understanding the roles of mutations and promoting factors in cancer. Eur J Epidemiol 38, 1251–1258 (2023).

6. Guerra, C. et al. Chronic Pancreatitis Is Essential for Induction of Pancreatic Ductal Adenocarcinoma by K-Ras Oncogenes in Adult Mice. Cancer Cell 11, 291– 302 (2007).

7. Bailleul, B. et al. Skin hyperkeratosis and papilloma formation in transgenic mice expressing a ras oncogene from a suprabasal keratin promoter. Cell 62, 697–708 (1990).

8. Brown, K. et al. v-ras genes from Harvey and BALB murine sarcoma viruses can act as initiators of two-stage mouse skin carcinogenesis. Cell 46, 447–56 (1986).

9. Elliot, A., Myllymäki, H. & Feng, Y. Inflammatory Responses during Tumour Initiation: From Zebrafish Transgenic Models of Cancer to Evidence from Mouse and Man. Cells vol. 9 1018 Preprint at 10.3390/cells9041018 (2020).

10. Moore, R. J. et al. Mice deficient in tumor necrosis factor-α are resistant to skin carcinogenesis. Nature Medicine 1999 5:7 5, 828–831 (1999).

11. Noy, R. & Pollard, J. W. Tumor-associated macrophages: from mechanisms to therapy. (2014) doi:10.1016/j.immuni.2014.06.010.

12. Coffelt, S. B., Wellenstein, M. D. & de Visser, K. E. Neutrophils in cancer: neutral no more. Nat Rev Cancer 16, 431–446 (2016).

13. Feng, Y., Santoriello, C., Mione, M., Hurlstone, A. & Martin, P. Live Imaging of Innate Immune Cell Sensing of Transformed Cells in Zebrafish Larvae: Parallels between Tumor Initiation and Wound Inflammation. PLoS Biol 8, e1000562 (2010).

14. Yan, C., Huo, X., Wang, S., Feng, Y. & Gong, Z. Stimulation of hepatocarcinogenesis by neutrophils upon induction of oncogenic kras expression in transgenic zebrafish. J Hepatol 63, 420–8 (2015).

15. Hanahan, D. Hallmarks of Cancer: New Dimensions. Cancer Discov 12, 31–46 (2022).

16. Colom, B. et al. Spatial competition shapes the dynamic mutational landscape of normal esophageal epithelium. Nat Genet 52, 604–614 (2020).

17. Murai, K. et al. Epidermal Tissue Adapts to Restrain Progenitors Carrying Clonal p53 Mutations. Cell Stem Cell 23, 687–699.e8 (2018).

18. Vermeulen, L. et al. Defining stem cell dynamics in models of intestinal tumor initiation. Science 342, 995–8 (2013).

19. Becker, W. R. et al. Single-cell analyses define a continuum of cell state and composition changes in the malignant transformation of polyps to colorectal cancer. Nat Genet 54, 985–995 (2022).

20. Marjanovic, N. D. et al. Emergence of a High-Plasticity Cell State during Lung Cancer Evolution. Cancer Cell 38, 229–246.e13 (2020).

21. LaFave, L. M. et al. Epigenomic State Transitions Characterize Tumor Progression in Mouse Lung Adenocarcinoma. Cancer Cell 38, 212–228.e13 (2020).

22. Pickering, C. R. et al. Mutational landscape of aggressive cutaneous squamous cell carcinoma. Clin Cancer Res 20, 6582 (2014).

23. Balmain, A. & Pragnell, I. B. Mouse skin carcinomas induced in vivo by chemical carcinogens have a transforming Harvey-ras oncogene. Nature 303, 72–74 (1983).

24. Myllymäki, H. et al. Metabolic Alterations in Preneoplastic Development Revealed by Untargeted Metabolomic Analysis. Front Cell Dev Biol 9, 684036 (2021).

25. Lee, R. T. H., Asharani, P. V. & Carney, T. J. Basal Keratinocytes Contribute to All Strata of the Adult Zebrafish Epidermis. PLoS One 9, e84858 (2014).

26. Akitake, C. M., Macurak, M., Halpern, M. E. & Goll, M. G. Transgenerational analysis of transcriptional silencing in zebrafish. Dev Biol 352, 191–201 (2011).

27. Ramezani, T., Laux, D. W., Bravo, I. R., Tada, M. & Feng, Y. Live Imaging of Innate Immune and Preneoplastic Cell Interactions Using an Inducible Gal4/UAS Expression System in Larval Zebrafish Skin. Journal of Visualized Experiments e52107 (2015) doi:10.3791/52107.

28. Lavoie, H., Gagnon, J. & Therrien, M. ERK signalling: a master regulator of cell behaviour, life and fate. Nature Reviews Molecular Cell Biology 2020 21:10 21, 607–632 (2020).

29. Zhao, X., Jiang, M. & Wang, Z. TPM4 promotes cell migration by modulating F-actin formation in lung cancer. Onco Targets Ther 12, 4055–4063 (2019).

30. Liu, J., Zhang, Y., Li, Q. & Wang, Y. Transgelins: Cytoskeletal Associated Proteins Implicated in the Metastasis of Colorectal Cancer. Front Cell Dev Biol 8, 573859 (2020).

31. Staquicini, D. I. et al. Intracellular targeting of annexin A2 inhibits tumor cell adhesion, migration, and in vivo grafting. Scientific Reports 2017 7:1 7, 1–11 (2017).

32. Hu, Y., Song, Z., Chen, J. & Caulin, C. Overexpression of MYB in the Skin Induces Alopecia and Epidermal Hyperplasia. Journal of Investigative Dermatology 140, 1204–1213.e5 (2020).

33. Hou, Y. et al. Cellular diversity of the regenerating caudal fin. Sci Adv 6, (2020).

34. Cokus, S. J. et al. Tissue-Specific Transcriptomes Reveal Gene Expression Trajectories in Two Maturing Skin Epithelial Layers in Zebrafish Embryos. G3 (Bethesda) 9, 3439–3452 (2019).

35. Blanpain, C. & Fuchs, E. Epidermal homeostasis: a balancing act of stem cells in the skin. Nat Rev Mol Cell Biol 10, 207 (2009).

36. Bressan, R. B. & Pollard, S. M. Transcriptional and epigenetic regulatory mechanisms in glioblastoma stem cells. Stem Cell Epigenet 231–255 (2020) doi:10.1016/B978-0-12-814085-7.00010-6.

37. Saghafinia, S. et al. Cancer cells retrace a stepwise differentiation program during malignant progression. Cancer Discov 11, 2638 (2021).

38. Spike, B. T. et al. A mammary stem cell population identified and characterized in late embryogenesis reveals similarities to human breast cancer. Cell Stem Cell 10, 183 (2012).

39. Wagner, D. E. et al. Single-cell mapping of gene expression landscapes and lineage in the zebrafish embryo. Science 360, 981 (2018).

40. Santos-Pereira, J. M., Gallardo-Fuentes, L., Neto, A., Acemel, R. D. & Tena, J. J. Pioneer and repressive functions of p63 during zebrafish embryonic ectoderm specification. Nature Communications 2019 10:1 10, 1–13 (2019).

41. Koster, M. I. & Roop, D. R. Mechanisms Regulating Epithelial Stratification. 10.1146/annurev.cellbio.23.090506.123357 **23**, 93–113 (2007).

42. Perdigoto, C. N., Valdes, V. J., Bardot, E. S. & Ezhkova, E. Epigenetic Regulation of Epidermal Differentiation. Cold Spring Harb Perspect Med 4, (2014).

43. Bao, X. et al. ACTL6a enforces the epidermal progenitor state by suppressing SWI/SNF-dependent induction of KLF4. Cell Stem Cell 12, 193–203 (2013).

44. Mulder, K. W. et al. Diverse epigenetic strategies interact to control epidermal differentiation. Nature Cell Biology 2012 14:7 14, 753–763 (2012).

45. Pastushenko, I. & Blanpain, C. EMT Transition States during Tumor Progression and Metastasis. Trends Cell Biol 29, 212–226 (2019).

46. Liu, S. et al. The Role of CD276 in Cancers. Front Oncol 11, (2021).

47. Vences-Catalán, F. et al. Targeting the tetraspanin CD81 reduces cancer invasion and metastasis. Proc Natl Acad Sci U S A 118, e2018961118 (2021).

48. Chu, W. et al. Neuropilin-1 promotes epithelial-to-mesenchymal transition by stimulating nuclear factor-kappa B and is associated with poor prognosis in human oral squamous cell carcinoma. PLoS One 9, (2014).

49. Sterneck, E., Poria, D. K. & Balamurugan, K. Slug and E-Cadherin: Stealth Accomplices? Front Mol Biosci 7, 138 (2020).

50. Subbalakshmi, A. R., Sahoo, S., Biswas, K. & Jolly, M. K. A Computational Systems Biology Approach Identifies SLUG as a Mediator of Partial Epithelial-Mesenchymal Transition (EMT). Cells Tissues Organs 211, 689–702 (2022).

51. Shibue, T. & Weinberg, R. A. EMT, CSCs, and drug resistance: the mechanistic link and clinical implications. Nature Reviews Clinical Oncology 2017 14:10 14, 611–629 (2017).

52. Zhou, H. M., Zhang, J. G., Zhang, X. & Li, Q. Targeting cancer stem cells for reversing therapy resistance: mechanism, signaling, and prospective agents. Signal Transduction and Targeted Therapy 2021 6:1 6, 1–17 (2021).

53. Yuan, S. et al. Ras drives malignancy through stem cell crosstalk with the microenvironment. Nature 2022 612:7940 612, 555–563 (2022).

54. Ji, A. L. et al. Multimodal Analysis of Composition and Spatial Architecture in Human Squamous Cell Carcinoma. Cell 182, 497–514 (2020).

55. Welch, J. D. et al. Single-Cell Multi-omic Integration Compares and Contrasts Features of Brain Cell Identity. Cell 177, 1873–1887.e17 (2019).

56. Powell, D., Lou, M., Becker, F. B. & Huttenlocher, A. Cxcr1 mediates recruitment of neutrophils and supports proliferation of tumor-initiating astrocytes in vivo. Sci Rep 8, 1–12 (2018).

57. Ries, A. et al. Activin A: an emerging target for improving cancer treatment? Expert Opin Ther Targets (2020) doi:10.1080/14728222.2020.1799350.

58. Batlle, E. & Massagué, J. Transforming Grown Factor-β Signaling in Immunity and Cancer. Immunity 50, 924 (2019).

59. Evrard, M. et al. Developmental Analysis of Bone Marrow Neutrophils Reveals Populations Specialized in Expansion, Trafficking, and Effector Functions. Immunity 48, 364–379.e8 (2018).

60. Manz, M. G. & Boettcher, S. Emergency granulopoiesis. Nature Reviews Immunology 2014 14:5 14, 302–314 (2014).

61. Hirai, H. et al. C/EBPbeta is required for ‘emergency’ granulopoiesis. Nat Immunol 7, 732–739 (2006).

62. Walters, K. B., Green, J. M., Surfus, J. C., Yoo, S. K. & Huttenlocher, A. Live imaging of neutrophil motility in a zebrafish model of WHIM syndrome. Blood 116, 2803–2811 (2010).

63. Soza-Ried, C., Hess, I., Netuschil, N., Schorpp, M. & Boehm, T. Essential role of c-myb in definitive hematopoiesis is evolutionarily conserved. Proc Natl Acad Sci U S A 107, 17304–17308 (2010).

64. Jin, H. et al. c-Myb acts in parallel and cooperatively with Cebp1 to regulate neutrophil maturation in zebrafish. Blood 128, 415–426 (2016).

65. Hyun, K. K., De La Luz Sierra, M., Williams, C. K., Gulino, A. V. & Tosato, G. G-CSF down-regulation of CXCR4 expression identified as a mechanism for mobilization of myeloid cells. Blood 108, 812 (2006).

66. Kowanetz, M. et al. Granulocyte-colony stimulating factor promotes lung metastasis through mobilization of Ly6G+Ly6C+ granulocytes. Proc Natl Acad Sci U S A 107, 21248–21255 (2010).

67. Hall, C. et al. Transgenic zebrafish reporter lines reveal conserved Toll-like receptor signaling potential in embryonic myeloid leukocytes and adult immune cell lineages. J Leukoc Biol 85, 751–765 (2009).

68. 68. Quail, D. F., et al. Cancer Focus: Neutrophil phenotypes and functions in cancer: A consensus statement. J Exp Med 219, 39 (2022).

69. Fridlender, Z. G. et al. Polarization of Tumor-Associated Neutrophil Phenotype by TGF-β: ‘N1’ versus ‘N2’ TAN. Cancer Cell 16, 183–194 (2009).

70. Mizuno, R., Kawada, K. & Sakai, Y. Prostaglandin E2/EP signaling in the tumor microenvironment of colorectal cancer. International Journal of Molecular Sciences vol. 20 Preprint at 10.3390/ijms20246254 (2019).

71. Gabrilovich, D. I., Ostrand-Rosenberg, S. & Bronte, V. Coordinated regulation of myeloid cells by tumours. Nature Reviews Immunology vol. 12 253–268 Preprint at 10.1038/nri3175 (2012).

72. Shaul, M. E. & Fridlender, Z. G. Cancer-related circulating and tumor-associated neutrophils – subtypes, sources and function. FEBS J 285, 4316–4342 (2018).

73. Hammond, F. R. et al. An arginase 2 promoter transgenic illuminates anti-inflammatory signalling in zebrafish. bioRxiv 2022.02.14.480079 (2022) doi:10.1101/2022.02.14.480079.

74. Feng, Y., Renshaw, S. & Martin, P. Live Imaging of Tumor Initiation in Zebrafish Larvae Reveals a Trophic Role for Leukocyte-Derived PGE2. Current Biology 22, 1253–1259 (2012).

75. Karras, P. et al. A cellular hierarchy in melanoma uncouples growth and metastasis. Nature 2022 610:7930 610, 190–198 (2022).

76. 76. van den Berg, M., et al. Active and opportunistic breaching of the basement membrane by immune cells during tumour initiation Developmental Cell. Under Submission (2018).

77. Torborg, S. R., Li, Z., Chan, J. E. & Tammela, T. Cellular and molecular mechanisms of plasticity in cancer. Trends Cancer 8, 735–746 (2022).

78. Oshimori, N., Oristian, D. & Fuchs, E. TGF-β Promotes Heterogeneity and Drug Resistance in Squamous Cell Carcinoma. Cell 160, 963–976 (2015).

79. Mascré, G. et al. Distinct contribution of stem and progenitor cells to epidermal maintenance. Nature 2012 489:7415 489, 257–262 (2012).

80. Rangel-Huerta, E. & Maldonado, E. Transit-Amplifying Cells in the Fast Lane from Stem Cells towards Differentiation. Stem Cells Int 2017, (2017).

81. Burdziak, C. et al. Epigenetic plasticity cooperates with cell-cell interactions to direct pancreatic tumorigenesis. Science (1979) 380, (2023).

82. Freisinger, C. M. & Huttenlocher, A. Live Imaging and Gene Expression Analysis in Zebrafish Identifies a Link between Neutrophils and Epithelial to Mesenchymal Transition. PLoS One 9, e112183 (2014).

83. Ng, M. S. F. et al. Deterministic reprogramming of neutrophils within tumors. Science (1979) 383, 1–16 (2024).

84. Campbell, J. S. et al. PTPN21/Pez Is a Novel and Evolutionarily Conserved Key Regulator of Inflammation In Vivo. Current Biology 31, 875–883.e5 (2021).

85. Westerfield, M. The Zebrafish Book: A Guide for the Laboratory Use of Zebrafish (Danio Rerio). (University of Oregon Press, 2000).

86. Myllymaki, H., Elliot, A. & Feng, Y. Wholemount Edu Staining (Zebrafish Larvae). (2023) doi:10.17504/PROTOCOLS.IO.N92LD9429G5B/V1.

87. Elliot, A. & Feng, Y. Zebrafish larvae dissociation for FACs sorting cells expressing fluorescent proteins. (2023) doi:10.17504/protocols.io.yxmvmk819g3p/v1.

88. Zebrafish larvae dissociation for FACs sorting cells expressing fluorescent proteins. https://www.protocols.io/view/zebrafish-larvae-dissociation-for-facs-sorting-cel-yxmvmk819g3p/v1.

89. Picelli, S. et al. Full-length RNA-seq from single cells using Smart-seq2. Nature Protocols 2013 9:1 9, 171–181 (2014).

90. Young, M. D. & Behjati, S. SoupX removes ambient RNA contamination from droplet-based single-cell RNA sequencing data. Gigascience 9, 1–10 (2020).

91. Wolock, S. L., Lopez, R. & Klein, A. M. Scrublet: Computational Identification of Cell Doublets in Single-Cell Transcriptomic Data. Cell Syst 8, 281–291.e9 (2019).

92. DePasquale, E. A. K. et al. DoubletDecon: Deconvoluting Doublets from Single-Cell RNA-Sequencing Data. Cell Rep 29, 1718–1727.e8 (2019).

93. Hafemeister, C. & Satija, R. Normalization and variance stabilization of single-cell RNA-seq data using regularized negative binomial regression. Genome Biol 20, 1–15 (2019).

94. Korsunsky, I. et al. Fast, sensitive and accurate integration of single-cell data with Harmony. Nature Methods 2019 16:12 16, 1289–1296 (2019).

95. Cao, J. et al. The single-cell transcriptional landscape of mammalian organogenesis. Nature 2019 566:7745 566, 496–502 (2019).

96. Liu, J. et al. Jointly defining cell types from multiple single-cell datasets using LIGER. Nature Protocols 2020 15:11 15, 3632–3662 (2020).

97. The Cancer Genome Atlas Program (TCGA) - NCI. https://www.cancer.gov/ccg/research/genome-sequencing/tcga.

98. Hobbs, G. A., Der, C. J. & Rossman, K. L. RAS isoforms and mutations in cancer at a glance. (2016) doi:10.1242/jcs.182873.

99. Hänzelmann, S., Castelo, R. & Guinney, J. GSVA: Gene set variation analysis for microarray and RNA-Seq data. BMC Bioinformatics 14, 1–15 (2013).

100. Pearce, D. A., Nirmal, A. J., Freeman, T. C. & Sims, A. H. Continuous Biomarker Assessment by Exhaustive Survival Analysis. bioRxiv 208660 (2018) doi:10.1101/208660.

